# No general effects of advancing male age on ejaculates: a meta-analysis across the animal kingdom

**DOI:** 10.1101/2023.04.14.536443

**Authors:** Krish Sanghvi, Regina Vega-Trejo, Shinichi Nakagawa, Samuel J.L. Gascoigne, Sheri Johnson, Roberto Salguero-Gómez, Tommaso Pizzari, Irem Sepil

**Author notes:** joint first authors (Krish Sanghvi, Regina Vega-Trejo).

## Abstract

Senescence, the deterioration of organismal function with advancing age, is a puzzling biological phenomenon. While actuarial senescence (*i.e.*, age-dependent increases in mortality rates) is well described across some taxa, reproductive senescence (*i.e.* age-dependent declines in reproduction) is less understood, especially in males, with mixed patterns reported across studies. To examine the evidence for male reproductive senescence, we investigated how advancing male age affects ejaculate traits across non-human animals via a meta-analysis yielding 1814 effect sizes from 379 studies. We found no evidence for a general pattern of reproductive senescence. Instead, we found high heterogeneity for how reproduction changes with male age across animals. Some of this heterogeneity (>10%) was associated with biological factors. For example, there were taxonomical differences for some ejaculate traits — sperm motility declined with male age in lab rodents and fish, whereas ejaculate size improved with male age in bulls, fish, and insects. Some methodological factors were also important in explaining this heterogeneity: studies sampling a larger proportion of a species’ lifespan were more likely to detect senescence in ejaculate traits, emphasising the need to examine the full life cycle of species to document senescence. Contrary to predictions, we reveal that the evidence for senescence in ejaculate traits is sporadic. Our findings will help generate novel hypotheses and identify more effective methodological approaches for studying male reproductive senescence.

## Introduction

Senescence is the age-dependent irreversible deterioration of organismal function, which leads to an increased risk of intrinsic mortality (Reinke et al, 2022) and decline in reproductive output (Lemaitre and Gaillard, 2017) with advancing age. Senescence has been reported in a number of taxa (Nussey et al, 2013), though it is not universal (Monaghan et al, 2008). Some studies have found negligible evidence for senescence (Jones et al, 2014; Roper et al, 2021), while others have found age-dependent improvements in organismal function (*i.e.* negative senescence: Jones and Vaupel, 2017; Vaupel at al, 2004). Additionally, the rate, pattern, and onset of senescence can differ greatly between individuals (Rodriguez-Munoz et al, 2019) and species (Jones et al, 2014; Nussey et al, 2013; Roper et al, 2021). Senescence is driven by mechanisms such as age-dependent deterioration of cellular repair, accumulation of mutations, oxidative damage, and telomere attrition in cells (Monaghan and Metcalfe, 2019). Senescence is also suggested to be caused by hyperfunctioning of cellular processes with increasing age, which leads to misactivation of signalling pathways (Blagosklonny, 2021). From an evolutionary perspective, senescence is hypothesized to be the result of relaxed selection against deleterious mutations in older organisms as first proposed by Medawar’s ‘mutation accumulation theory’ (Medawar, 1952; Lemaitre et al, 2015). Other proposed explanations for senescence include selection for alleles that increase performance early in life, but convey net costs later in life (‘antagonistic pleiotropy’), and trade-offs between investment in survival *versus* reproduction (‘disposable soma’: Charlesworth, 1993; Rose and Charlesworth, 1980; Kirkwood, 1977; Kirkwood and Austad, 2000; Kirkwood and Rose, 1991; Stearns, 1989; Williams, 1957).

Reproductive senescence has been relatively well documented in females (Archer et al, 2022; Ivimey-Cook et al, 2023; Nussey et al, 2013; Tarin et al, 2000). Yet, patterns, causes, and consequences of male reproductive senescence are much less understood (Fricke and Koppik, 2019; Kirkwood and Austad, 2000; Lemaitre and Gaillard, 2017). This gap of knowledge is perhaps due to challenges in assigning paternity and measuring reproductive success in males. Understanding male reproductive senescence is crucial for several reasons. First, compared to females, males often display greater variance in reproductive success because of more intense intrasexual competition (Parker and Pizzari, 2010), potentially leading to stronger selection on male reproductive performance than that of females (Parker 1970, 1982, 2006). Thus, age-dependent changes in male ejaculate traits such as sperm number and motility, which underpin male reproductive performance, can modulate the intensity of sexual selection. For instance, senescence in male ejaculate traits can influence the outcome of sperm competition (Aich et al, 2021; Gasparini et al, 2010), affect cryptic female choice (Jones, 2002), and lead to sexual conflict (Adler and Bondurianski, 2014; Bondurianski et al, 2008; Dean et al, 2010; Parker, 2006; Radwan, 2003). Second, sperm are potentially more affected by advancing organismal age than eggs (Bronikowski et al, 2022; Nussey et al, 2009). This pattern is due to sperm being more active and shorter lived than eggs, and male germlines having higher rates of cell divisions and mutations (Bergeron et al. 2023; Crow, 2000; Fischer and Riddle, 2018; Monaghan and Metcalfe, 2019), but poorer DNA repair machinery, than female germ cells (Reinhardt and Turnell, 2019). Consequently, there may be greater transmission of accumulated mutations to offspring, via the paternal germline (Bergeron et al. 2023; Campbell and Eichler, 2013; de Manuel et al, 2022; Sharma et al, 2015). Third, male reproductive senescence can affect male and female fitness due to declines in fertilisation ability of males (Johnson and Gemmell, 2012; Paul and Robaire, 2013), or due to declines in offspring quality via paternal age effects (Eisenberg and Kuzawa, 2018; Pizzari et al, 2008; Schroeder et al, 2015; Priest et al, 2002; Preston et al, 2015). Lastly, males are hypothesized to invest more than females in early-life reproduction and follow a “live fast, die young” strategy, which may expose them to faster rates of senescence later in adult life than females (Adler and Bonduriansky, 2014; Bondurainsky et al, 2008; Vinogradov, 1998).

Several studies have shown that compared to younger males, older males have poorer ejaculate traits that influence male reproductive success and offspring quality. Specifically, senescence has been observed in: ejaculate volume (de Fraitpont et al, 1993), mitochondrial function (Wei et al, 2019), sperm concentration (Sasson et al, 2012), motility (Johnson et al, 2018), viability (Gasparini et al, 2010), velocity (Aich et al, 2021; Dean et al, 2010; Gasparini et al, 2019), number (Cornwallis et al, 2014), telomere length (de Frutos et al, 2016), and morphological (Delbarco-Trillo et al, 2018), DNA (Velando et al, 2011), and oxidative damage (Selvaratnam and Robaire, 2016). Other studies however, have reported age-dependent improvement in ejaculate traits. For instance, younger males having poorer ejaculate traits (e.g. Decanini et al, 2013; Heinze et al, 2018; Lifjeld et al, 2022), smaller ejaculates (e.g. Aich et al, 2021; Kehl et al, 2015; Vega-Trejo et al, 2019; Wedell and Ritchie, 2004), and lower ferilisation success (Lifjeld et al, 2022) than older males. There are theoretical reasons for why advancing male age might lead to an improvement in male reproductive output, instead of net deterioration (*i.e.* senescence; Johnson and Gemmell, 2012). For example, the “good genes” hypothesis predicts that on average, old males may have better reproductive performance than young males, because older cohorts are purged of lower quality phenotypes due to viability selection (Brooks and Kemp, 2001; Segami et al, 2021). Finally, some studies report no net changes in ejaculate traits with advancing male age (e.g. Delbarco-Trillo et al, 2018; Mickova et al, 2023).

Heterogeneity in the observed effects of male age on ejaculate traits may additionally be caused by various biological and methodological differences between studies (Johnson et al, 2015; Fricke and Koppik, 2019; Ricklefs, 2008). Heterogeneity could be associated with studies being conducted on different taxa, from insects (e.g. Ruhmann et al, 2018) to mammals (e.g. Xu et al, 2014); or different populations, from wild (e.g. Girndt et al, 2019) to laboratory animals (e.g. Sepil et al, 2020). Heterogeneity could also be due to studies sampling males in different ways, such as repeated (longitudinal) sampling of the same individuals (e.g. Gasparini et al, 2019), or cross-sectional sampling of different males in different age cohorts (e.g. Aich et al, 2021). Studies also differ in how ejaculates are collected, from more “natural” methods such as mating males with females (e.g. Jones et al, 2007), to highly invasive methods such as dissection (e.g. Turnell and Reinhardt, 2020).

Additionally, some studies use experimental manipulations of male social and biotic environments, such as mating history (e.g. Vega-Trejo et al, 2019) or diet (e.g. Guo and Reinhardt, 2020), while others do not. These biological and methodological factors (“moderators”) can in principle modulate the effect of advancing male age on ejaculate traits (see Table 1). Given the diversity of approaches and heterogeneity in studies, a meta-analytical approach is essential to understand the general effects of advancing male age on ejaculate traits. Yet, no study has done this across non-human animals (see Johnson et al, 2015 for humans; Vrtilek et al, 2022 for a systematic review in fish; and Fricke et al, 2023 for effects of male age on seminal fluid).

**Table 1:** Possible influence of different biological (highlighted in yellow) and methodological (highlighted in blue) moderators on male reproductive senescence at the level of the ejaculate. Moderators marked with an asterisk were included in our meta-analysis, because there were sufficient data across studies.

| Biological moderators | Possible influence |
| --- | --- |
| Taxon-specific effects* | Phylogenetic history and biology specific to taxa, such as ecosystems, niches, metabolic rates, life-history strategies, mating systems, and social and ecological conditions, could influence how male age affects ejaculate traits (Jones et al, 2008; Jones et al, 2014; Reinke et al, 2022; Roper et al, 2021). |
| Ejaculate traits* | Evidence for reproductive senescence can depend on the specific trait measured (Naciri et al, 2022). This can be due to trade-offs between different ejaculate traits (Snook, 2005) or different traits being under different selection pressures (Hayward et al, 2015; Moorad and Ravindran, 2022). For instance, males may show higher sperm motility at the cost of lower sperm viability and greater oxidative damage to sperm; or higher sperm viability and velocity at the cost of producing fewer sperm (Levitan, 2000; Snook, 2005). |
| Degree of sperm competition* | Species with increased levels of sperm competition have evolved increased investment in competitive ejaculate traits such as sperm number and velocity (Immler et al, 2011; Lupold et al, 2020), which may reduce the rate of senescence in these ejaculate traits (Delbarco-Trillo et al, 2018). However, the influence of sperm competition on male reproductive senescence likely depends on the life-history of a species. For example, in some species, males may preferentially invest resources in sperm competition early in life, and thus suffer faster rates of reproductive senescence later in life (Lemaitre et al, 2020). |
| Shape of ageing* | Rather than a constant rate of decline, ageing patterns are often characterised by an initial period of maturation where performance increases from early- to mid-adult life, and subsequently decreases (*i.e.* senescence) in late-adult life (Baudisch, 2011; Baudisch and Stott, 2019; Jones et al, 2014; Keyfitz, 1972; Reinke et al, 2022; Wrycza et al, 2015). Such curvilinear patterns of ageing may lead to an |
|  | underestimation of senescence. For instance, if ejaculate traits peak at an intermediate age, and if older males are scarcer, the positive part of the function (young to intermediate age) will be easier to detect than the negative part of the function (intermediate to old age) (Pizzari et al, 2008). |
| Life-history strategies | Specific life-history strategies (schedules of survival and reproduction along the life cycle) of populations, and pace of life (coevolution between vital rates and environmental conditions) of individuals, can modulate the effects of male age on reproductive traits (Bonduriansky et al, 2008; Healy et al, 2019; Maklakov et al, 2009). Additionally, *K-* versus *r*-selected species may differ in their pace of life and patterns of reproductive senescence, due to differential investment in quantity versus quality of offspring, and in lifespan versus reproduction (Danko et al, 2018; Jones et al, 2008; Healy et al, 2018; Service et al, 1987; Salguero-Gomez et al, 2016; Vinogradov, 1998). |
| Seminal fluid changes | Seminal fluid, which is a cocktail of micro- and macro-molecules such as proteins, carbohydrates, and vitamins, makes up a major part of ejaculates and is produced by somatic tissue (Fricke and Koppik, 2019; Perry et al, 2013). Levels of antioxidants in seminal fluid (Mannucci, 2021) and abundance of seminal fluid proteins can change as males age (Ruhmann et al, 2018; Sepil et al, 2020), independent of changes in sperm. These age-dependent changes in seminal fluid can impact sperm motility, sperm usage, sperm activation, sperm oxidative and DNA damage (Avila et al, 2011; Poiani, 2006), over and above the direct effects of male age on sperm (Fricke et al, 2023). |
| Ontogeny of secondary sexual traits | The ontogeny of secondary sexual traits can influence the evolution of male reproductive senescence rates (Bonduriansky et al, 2008). For instance, in species where secondary sexually selected traits such as weapons or ornaments improve with age, males are hypothesized to evolve lower rates of reproductive senescence, compared to species where these male traits do not improve with age (Archer and Hunt; 2015; Bonduriansky et al, 2008; Graves, 2007; Lifjeld et al, 2022; Promislow, 2003). |
| Age-dependent mortality | Organisms that evolve in environments with high extrinsic mortality, may show faster rates of senescence when old, because deleterious late-life expressed alleles would not be selected against (Abrams, 1993; Moorad et al, 2019; Vaupel et al, 1979; Williams and Day, 2003). Similarly, animals may invest more in early-life reproduction when age-dependent mortality risk is high (Hechinger, 2010), and thus show higher senescence rates than animals facing lower age-dependent mortality risk (Danko et al, 2018; Nussey et al, 2013; Reznick et al, 2004; Ricklefs, 1998). |
| **Methodological moderators** | **Possible influence** |
| Manipulations * | Manipulated environments that are outside what healthy organisms typically experience, such as environments with stressful conditions, can exacerbate reproductive senescence (Cooper and Kruuk, 2018; Sanghvi et al, 2021, 2022). Thus, males exposed to manipulations such as thermal stress, poor diet, or toxins would be more likely to show reproductive senescence than males not subjected to these. Other manipulations, such as experimental inbreeding (Keller et al, 2008; Langen et al, 2017) or lines with deleterious mutations (Butterfield and Poon, 2005; Yousefzadeh et al, 2019), may exacerbate senescence. |
| Population type* | Senescence rates have been shown to differ between animals in captive *versus* wild populations (Kawasaki et al, 2008; Reichard, 2016; Williams et al, 2006; Zajitschek et al, 2020). Additionally, domestic animals are often culled prior to reaching the onset of senescence (Workie et al, 2021), and have undergone generations of artificial selection for unusual life-histories (e.g. extremely short generation time in broiler chicken; Gering et al, 2019; Tallentire et al, 2018). Studies on captive, domestic, wild, and lab populations may thus yield differing patterns of senescence. |
| Cross-sectional *versus* longitudinal male sampling* | Cross-sectional sampling of males makes individual-level reproductive senescence harder to detect (Nussey et al, 2008), especially if low-quality males selectively disappear (Bouwhuis et al, 2009; Hamalainen et al, 2014; Sanghvi et al, 2022). This is because non-random age-dependent mortality of poor-quality males can lead to a biased sampling of old males, where younger age classes would have higher variance but lower means in ejaculate traits than old age classes. Thus, cross-sectional studies might underestimate male reproductive senescence, compared to longitudinal sampling measuring the same individuals at different ages. |
| Ejaculate collection method* | Ejaculates are collected in different ways (e.g. electroejaculation, masturbation, natural mating). If a male has control over ejaculation (e.g. natural mating), males may have the opportunity to strategically adjust ejaculate phenotypes (Kelly and Jennions, 2011; Wedell et al, 2002), causing age-independent changes and reducing the detectability of senescence in ejaculate traits. Additionally, when males have control over ejaculation, studies may obtain a smaller proportion of the sperm reserves available to a male, which may not be representative of a male's whole-ejaculate phenotype, compared to studies that use invasive methods and obtain the whole-ejaculate (e.g. dissection). However, invasive methods may capture cells (e.g. immature sperm) that do not typically contribute to a natural ejaculate. |
| Proportion lifespan sampled* | A higher proportion of lifespan sampled will increase the probability of detecting reproductive senescence, as the onset of senescence usually occurs late in life (Gaillard and Lemaitre, 2020; Fricke et al, 2023, Jones et al, 2014; Lemaitre et al, 2020; Peron et al, 2010). |
| Mating history and post-meiotic sperm storage | Temporal changes in sperm traits can occur not just with advancing male age, but also due to post-meiotic storage of mature sperm in males before ejaculation, and in females following mating (Pizzari et al, 2008; Vega-Trejo et al, 2019). The duration since last mating of males (*i.e.* sexual rest) can influence the amount of post-meiotic damage to sperm, such that for a given age, males with shorter sexual rest (e.g. high mating rate) will incur lower post-meiotic sperm damage (Jones and Elgar, 2004; Siva-Jothy, 2000). However, males with long sexual rest durations (e.g. virgins) may accumulate more sperm and thus produce larger ejaculates (Sepil et al, 2020). |

Here, we conduct a systematic review and meta-analysis to address three aims. First, we test how advancing male age affects ejaculate traits (*aim 1*). We predict, based on the most common theories of ageing (mutation accumulation, disposable soma, antagonistic pleiotropy), that senescence in ejaculate traits will be observed consistently across the examined animal species. Second, we investigate the role of biological and methodological moderators in modulating the effects of male age on ejaculate traits (*aim 2*). Finally, we quantify how advancing male age affects reproductive traits, such as male fertilisation success and fecundity, and whether effects of male age on ejaculate traits differ from effects on reproductive traits (*aim 3*). Although various possible moderators can influence the effects of advaing male age (Table 1), we selected moderators (indicated by an asterisk in Table 1; most were pre-registered on OSF https://doi.org/10.17605/OSF.IO/2D8Z6) that are commonly reported across most studies to test their influence on modulating effects of male age on ejaculate traits.

## Materials and methods

We conducted a meta-analysis following the PRISMA-EcoEvo guidelines (O’Dea et. al. 2021), with all statistical analyses conducted in R v 4.1.2 (R core Team, 2022). The data, code, and complete model outputs will be available upon acceptance, and supplementary figures are presented in the “Supplementary figures” file.

### Search protocol

We first conducted a scoping search on Google Scholar. Our scoping search was done using the keywords “male age sperm -human”, and we used the first 48 papers (*i.e.* all papers from the first 5 pages out of 11800 pages of the search) to build a word cloud to discern the most common words used in these studies. We used the most commonly occurring relevant words from this word cloud to create keywords for our search string (Appendix 1). We conducted a literature search using this search string on SCOPUS and Web of Science (WoS), accessed through the University of Oxford server on 21^st^ January and 27^th^ March 2021, respectively (See Appendix 2 for specific search strings based on a scoping search). In addition, we conducted a backwards- and forwards- search using seven papers related to the topic of our meta-analysis. Four of these papers were the most cited reviews (Fricke & Koppik, 2019; Monaghan and Metcalfe, 2019; Pizzari et al, 2008; Radwan, 2003), while three were the most cited research articles (standardized by unit time) on the topic of pre-meiotic sperm senescence in the fields of ecology and evolution in the last 20 years (Gasparini et al, 2010; Jones and Elgar, 2004; Vega-Trejo et al, 2019). To avoid a bias towards peer-reviewed published literature, we additionally conducted a search for unpublished Ph.D. and M.Sc. theses using the database BASE (Bielefield Academy Search Engine; Pieper and Summann, 2006). Finally, we contacted 56 researchers who study the ecology and evolution of male reproductive senescence to ask for unpublished data.

Our search resulted in 7026 hits from SCOPUS (1923- 2021) and 4035 hits from WoS (1951 - 2021). We further obtained 959 papers from the back- and forwards- search, an additional 21 papers from other sources (*i.e.* a previously collected dataset provided by SN and SJ), unpublished data from 1 researcher, and 271 Ph.D. theses from BASE (Prisma diagram in Appendix 3 and Supplementary Fig. 1). Our search resulted in a total of 9412 unique abstracts from published sources, and 271 abstracts from unpublished sources, which we screened in Rayyan (Ouzzani et al, 2016) and abstrackr (Rathbone et al, 2015) using pre- defined inclusion and exclusion criteria (see below). KS screened all abstracts to check the suitability of each study (n = 9683) while SG checked for repeatability by screening ∼50% of abstracts (n = 4918). KS and SG agreed on the inclusion or exclusion (*i.e.* suitability) of 91% of the studies (Kappa = 0.56; Kappa calculates inter-rater reliability for qualitative items; Koricheva et al, 2013) when screening abstracts. KS then screened all full texts that passed the abstract screening stage (n = 1003 studies from published sources and four from unpublished sources) to determine whether a study was suitable to be included in the meta-analysis, while SG screened ∼10% of full texts to test for repeatability (n = 100). KS and SG agreed on the inclusion or exclusion of 98% of the studies when screening full-texts (Kappa = 0.96).

### Inclusion criteria

For a study to be included in our full-text screening, we imposed further selection criteria. Specifically, the study had to be a research article (not a review or meta-analysis) written in English, on non-human animals, and quantify ejaculate traits in males of different ages. Additionally, the study needed to include data for more than a single male per age group (*i.e.* it could not be a case study). The study also needed to contain: data on effects of male age on ejaculate traits, non-overlapping age groups of males, and appropriate data for calculation of effect sizes. We only included studies where different ages of adult males could be compared. Thus, if all age groups in a study had immature (non-adult) males, we excluded these studies. On the other hand, if at least two age groups of males were adults, we included the study, and we used only the data from adult males. Groups that were described as “[pre-] pubertal”, “adolescent”, “juvenile”, or “immature” were not considered as adults and thus not included in the analysis, because we were only interested in age-dependent changes in adult individuals. For arthropods, we included all post-eclosion/last-moult ages in the analysis because this is when arthropods are usually considered adults (Rolff et al, 2019). Whenever data on means, SE/SD, or sample sizes were not reported clearly in the study, we contacted the authors via a standard email to request these data, and the study was excluded if the authors could not help or did not respond. We deemed a total of 379 studies (374 from published, and five from unpublished sources) appropriate for data extraction based on our inclusion criteria, and included them in our meta-analysis.

### Data collection

#### Information for calculation of effect sizes

To quantify the evidence for or against male reprodcuctive senescence (*aim 1*), we collected quantitative data. These data were: means, standard deviations or standard errors, the number of males in each age group, and the number of unique males in the study, wherever reported. We converted standard errors to standard deviations (SD) using the formula SD = SE* sqrt (N), to calculate effect sizes (Nakagawa and Cuthill, 2007). When medians and inter-quartile ranges were reported, we converted these to SD following Wan et al (2014). We collected data (in the following order) from: tables, text in the Results section, figures using WebPlotDigitizer (Rohtagi, 2014) and MetaDigitize (Pick et al, 2019), the supplementary information or raw data provided with the paper, a data repository, or by directly contacting the authors of the paper. If we could not obtain means and SD/SE using these methods, we noted the “test statistic” reported in the study from which effect sizes can be easily obtained (formulae from Koricheva et al, 2013; Polanin and Stiltsveit, 2016). These test statistics included: correlation coefficients (Pearson’s or Spearman’s), R squared or adjusted R squared values, F statistics from ANOVA and ANCOVA with 1 DF, z values from z-tests, U from Mann-Whitney U tests, and T from Seigel’s T or t-tests with 1 DF.

To assure repeatability of data extraction, we checked data obtained from ∼5% of papers (*i.e.* 19 studies and 75 effect sizes). For this, two people, RVT and KS first independently extracted data and calculated effect sizes (Fisher’s z-transformed correlation coefficient) for each of the 75 rows of the data collected. Then a coefficient of determination between the effect sizes obtained by RVT and KS was calculated (R^2^ = 0.96, P<0.001), indicating strong repeatability.

##### Biological moderators

To understand how biological moderators affect patterns of senescence (*aim 2*), we recorded various biological variables from the 379 studies included in the meta-analysis. These variables were: the species and taxonomic class of the study organism; whether males belonged to wild, domestic, captive or laboratory populations (See Appendix 4 for definitions); the maximum lifespan and age at adulthood of the species studied (See Appendix 5). We then calculated the proportion of maximum adult lifespan sampled for a species in each study (converted to years). We additionally recorded the level of sperm competition (measured as gonadosomatic index) for the species reported in the study, wherever possible (See Appendix 5). Finally, we recorded the ejaculate traits measured in the study (See Appendix 6 for definitions of traits). The traits were either measures of sperm/ejaculate quantity (e.g. sperm concentration, sperm number, and ejaculate volume), sperm performance (e.g. sperm motility, velocity, viability), or intra-cellular measures of sperm quality (e.g. oxidative stress in sperm, DNA damage to sperm, sperm telomere length).

#### Methodological moderators

To test how methodological moderators affect the patterns of senescence in ejaculate traits (*aim 2*), we collected data on various methodological variables from included studies (see Table 1). Specifically, we quantified: whether males were sampled longitudinally or not (*i.e.* the same males repeatedly measured at different ages, or males from different age cohorts measured once); method of sperm extraction (e.g. electro-ejaculation, natural mating); method for measuring male age (*i.e.* whether male age was known directly or estimated from a measure of phenotype indirectly); whether the ejaculate was stored in cold conditions (<5°C, irrespective of the duration of storage) before analysis of sperm velocity, motility, and viability; and whether the study was experimental or not (non-experimental studies did not manipulate something in addition to male age, and opportunistically sampled the available age class distributions: following Pinquart et al, 2022; Vrtilek et al, 2022). In studies where males underwent “unnatural manipulations” (See Appendix 7 for detailed definitions), we also recorded whether the data were obtained from males that underwent these “unnatural” manipulations (*i.e.* males that experienced conditions outside of their typical range and that were compared to a well-defined control in the study), or from males that were used as controls in the same study.

#### Reproductive traits

To test whether male age-driven changes affect male reproductive traits (*aim 3*), and whether these effects of male age on reproductive traits are different from the effects on ejaculate traits, we collected data on various reproductive traits from included studies. Specifically, we collected data on how advancing male age affects: male fertilization success, number of eggs produced by the mated females, number of offspring produced by the mated females, egg viability, offspring viability, offspring developmental rate, and offspring body condition, whenever available (53 studies; See Appendix 6).

### Calculating effect sizes

We used Fisher’s z transformed correlation coefficient (henceforth referred to as Zr) as the effect size in our meta-analysis. We calculated Zr values from correlation coefficients (r) using the formula: Zr = 0.5*(log(1+ r) - log(1-r)) (Nakagawa and Cuthill, 2007). Because r values were rarely ever directly reported in studies, we had to calculate these indirectly. For studies with two age groups, we calculated correlation coefficients (r) using standardized mean differences (SMD). When there were more than two age groups, we calculated correlation coefficients using a simulation (See Appendix 9). For studies where only test statistics were reported, we calculated correlation coefficients using test-specific formulae (See Appendix 9). We corrected all calculated effect sizes (Zr) for a multiplier (See Appendix 10) to obtain a final effect size to be used in the analyses. Negative effect sizes indicated older males having worse sperm and ejaculate traits (*i.e.* senescence) than younger males, while positive effect sizes indicated an improvement in these traits with advancing age. To ensure that the three different effect size calculation methods did not affect the overall outcome in our meta-analysis, we compared the meta-analytical mean obtained from each method (*i.e.* SMD, simulation, and test-statistics). These measures of effect sizes did not differ from each other (Appendix 11), hence we analysed effect sizes calculated from SMD, simulation, and test-statistics together in subsequent models.

### Data analysis

#### Aim 1: Effects of advancing male age on ejaculate traits

##### Meta-analytical model

We first created a meta-analytical model (*i.e.* “null” model) to test for the general overall effect of advancing male age on ejaculate traits. We included the effect size (Zr) as our response variable in the null model. This null model included random effects of: effect size ID, cohort ID, study ID, and species name to control for non-independence of effect sizes (Noble et al, 2017). We also added a correlation matrix related to the phylogenetic relatedness of species in our dataset, to control for non-independence arising due to shared phylogenetic history and test for a phylogenetic signal (Cinar et al, 2021; Nakagawa and Santos, 2012; Noble et al, 2017). The phylogenetic tree (Supplementary Fig. 2) was built using the packages *ape* (Paradis et al, 2004) and *rotl* (Michonneau et al, 2016), which use data from the OpenTreeOfLife (Hinchliff et al, 2015). We quantified the total heterogeneity not due to sampling error as 𝐼^2^(Gurevitch et al, 2018; Higgins and Thompson, 2002), which can range from 0-100. Next, we quantified partial heterogeneity explained by each random effect using the function *i2_ml* from the *orchard* package (Nakagawa et al, 2020). We built the meta-analytical and meta-regression models using *rma.mv* in *metafor* (Viechtbauer, 2010).

#### Aim 2: Role of biological and methodological moderators

##### Meta–regressions

We created meta-regressions to test how various moderators influenced the effects of advancing male age. In all meta-regressions, we included the same random effects and phylogenetic matrix as our null model, and effect size (Zr) as our response variable. For each meta-regression model, we calculated the proportion of heterogeneity explained by moderators (marginal 𝑅^2^) with the function *r2_ml,* and the total heterogeneity explained (QM), using the *orchard* package. P values (α = 0.05) indicate whether the heterogeneity explained was significant or not (Higgins and Tompson, 2002). We created models without an intercept (*i.e.* each level of the moderator was compared against zero) to test whether each level/category of a moderator showed evidence for senescence or improvement of ejaculate traits with age. For some moderators with two levels, we were additionally interested in testing whether effect sizes in one level of the moderator were significantly different from those in the other level. In such cases, we created a model with one level of the moderator as the intercept (here, a P value expressed whether one level of the moderator was different from the other level).

We first conducted a meta regression with all moderators for which data were available for >75% of effect sizes and studies (“full” model). This full model was constructed to estimate the proportion of heterogeneity explained by moderators (Noble et al, 2022; Senior et al, 2016), while accounting for the confounding effects of other moderators. This full model included moderators of: taxonomic class, ejaculate trait, population type, proportion of maximum adult lifespan sampled, whether or not males had control over ejaculation, method of age estimation, sampling method, whether or not a study was experimental, and whether or not males underwent “unnatural” manipulations. To test for robustness of our full and null models, we created a variance-covariance matrix (VCV) of correlation values between effect size ID and cohort ID that replaced the variance argument (following Nakagawa et al, 2021) in both, null and full models. The results obtained for the null model or full model were not different between the model that used the VCV matrix *versus* the one that did not, thus we present the model without the VCV matrix throughout.

We then built several meta-regressions to explore individually the effects of each methodological and biological moderator, in line with our hypotheses (See Table 1, Appendix 8, and pre-registration). Here, we also tested two additional moderators that were not pre-registered and had data for <75% of studies and effect sizes: gonadosomatic index of species (GSI, used as a proxy for sperm competition; Parker et al, 2017; Stockley et al, 1997) and whether ejaculates underwent cold storage before analysis of sperm performance. The individual moderator models do not control for possible confounding effects of other moderators, and should be seen as hypothesis generating, rather than demonstrating causality (Baker et al, 2009).

Some taxonomic groups were over-represented in our dataset. Insecta, Actinopterygii, Aves, and Mammalia were over-represented classes in our dataset, as they included more than 150 effect sizes each (out of a total of 1814 effect sizes) from more than 30 studies each (Supplementary Fig. 2). We thus created a model where we included an interaction between class and ejaculate trait (by sub-setting these four classes), as well as four separate meta-regressions to test for how ejaculate traits change with age in each of these four classes. For the four most frequently occurring species in our dataset*, i.e.* lab mice (*Mus musculus*) and lab rats (*Rattus norvegicus*) combined, chicken/red junglefowl (*Gallus spp.*), and bulls (*Bos taurus*) (each species being represented by >150 effect sizes across >20 studies, Supplementary Fig. 2), we further created three separate meta-regression models with ejaculate trait as a moderator.

#### Aim 3: Effects of advancing male age on reproductive traits

To compare age-dependent changes in ejaculate traits *versus* male reproductive traits, we used data from included studies that measured age-dependent changes in both ejaculate traits and reproductive traits (see Appendix 6 for definitions). Then we ran a meta-regression using type of trait (reproductive trait vs ejaculate trait) as a moderator.

### Other analyses

#### Quadratic effects of age

Our meta-analyses used an effect size (Zr), which assumes a linear relationship between independent (age) and dependent (ejaculate traits) variables (Baker et al, 2009). Thus, to test whether the effects of advancing male age on ejaculate traits are curvilinear in shape, we created linear mixed models (LMM) using the package *lme4* (Bates et al, 2014) and *lmerTest* (Kuznetsova et al, 2017). We standardized each of the different ages at which males were sampled at, as the proportion of maximum adult lifespan of the species (independent variable). We standardized the traits (response variable) by calculating the percentage of: morphologically normal sperm (85 studies), viable sperm (137 studies), and motile sperm (81 studies), and analyzed them in three separate models. We included linear and quadratic effects of [standardized] age (covariates), with population type (i.e, lab, captive, domestic, wild) and taxonomic class as fixed effects. We also used the sample size of males within each age class (log transformed) as weights in the LMM. We included effect size ID, cohort ID, and study ID as random effects. Our models met assumptions of homoscedasticity and normality of residuals, checked using the *stats* package (R Core Team, 2022).

#### Publication bias

We tested for publication bias using various methods (following Nakagawa and Santos, 2012; Nakagawa et al; 2022). We first visually evaluated symmetry in a funnel plot with effect sizes (Zr) on the X axis, and the inverse of the standard error (1/SE) of the effect size (*i.e.* precision) on the Y axis (note that for Zr, precision is proportional to sample sizes). Second, we conducted a multi-level meta-regression to evaluate whether the size of a study and its year of publication influences effect sizes (*i.e.* small study bias and time-lag bias) by including both, standard error of effect sizes and year of publication as moderators, and effect size ID, cohort ID, study ID, species name, and phylogeny as random effects. Third, we created a funnel plot with the average of effect sizes from each study (X axis) against precision (Y axis), and tested for funnel asymmetry using a trim-and-fill method (Nakagawa et al, 2022). Finally, we created a selection model to test whether the probability of selecting a study depended on the significance of its effect size (Marks-Anglin and Chen, 2020).

## Results

### Aim 1: Effects of advancing male age on ejaculate traits

We found no general effect of advancing male age on ejaculate traits (mean [95% confidence interval]: -0.006 [-0.486 to 0.474], z = -0.025, P = 0.978, Fig. 1A). Heterogeneity in our dataset was high (𝐼^2^= 95 %), with 40% attributed to true differences between studies, 19% to observation-level differences (*i.e.*, effect size ID), 0.6% to differences between cohorts, and 0% to between-species differences. Phylogenetic relatedness (Supplementary Fig. 2) explained ∼35% of variance in effect sizes, suggesting a phylogenetic signal on male reproductive senescence.

**Figure 1:**
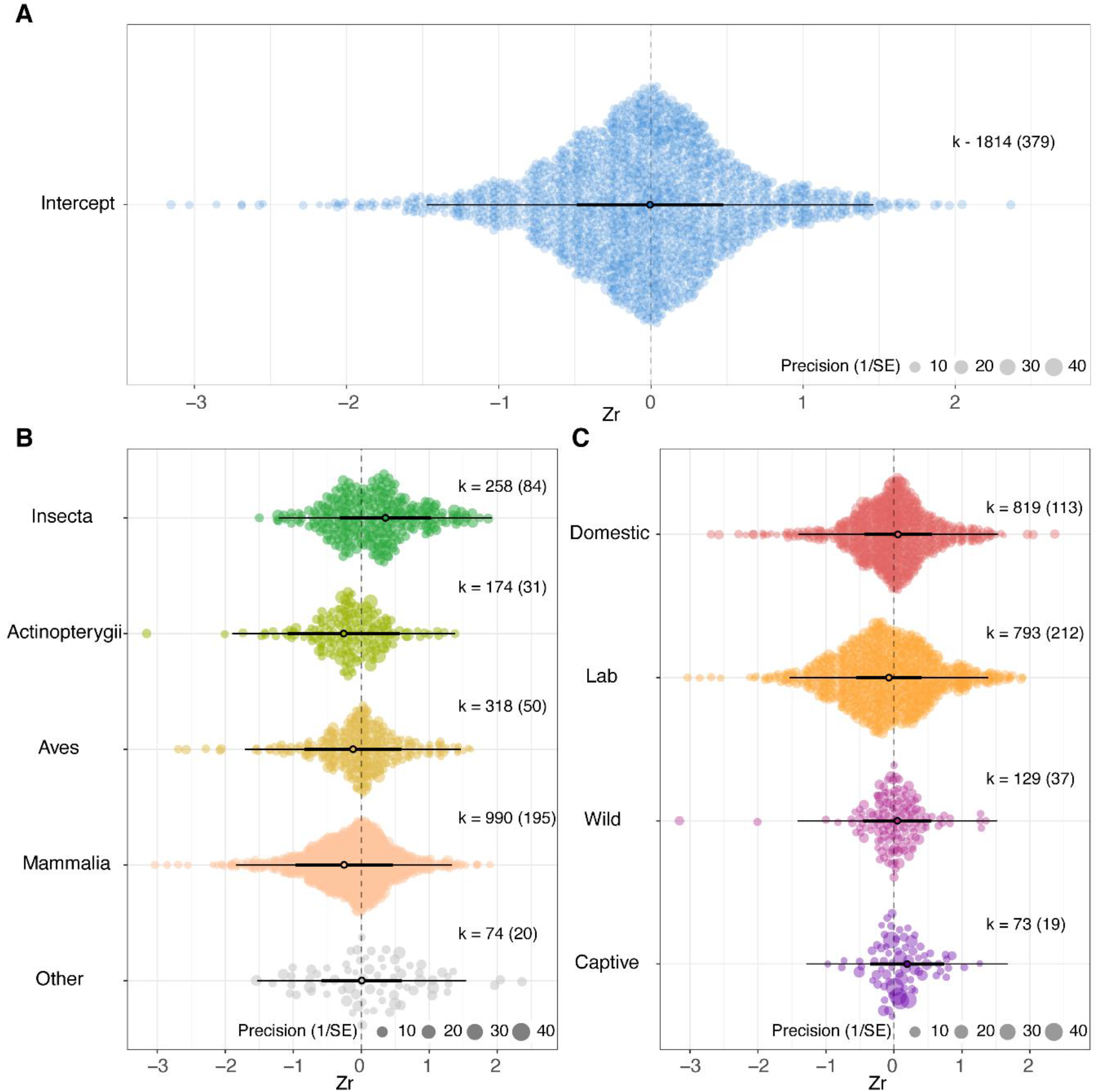
No consistent evidence for senescence in ejaculate traits, irrespective of the study population type or taxonomic class. **A.** Meta-analytical model of the overall effect of advancing male age on ejaculate traits. **B.** Effect of advancing male age on ejaculate traits for each taxonomic class (note that animal classes with less than 25 effect sizes were grouped together in ‘Other’). **C.** Effect of advancing male age on ejaculate traits for different types of population. The size of each data point represents the precision of the effect size (1/SE). The X axis represents values of effect sizes as Fisher’s z-transformed correlation coefficient (Zr), while the Y axis shows the density distribution of effect sizes. The position of the overall effect is shown by the dark circle, with negative values depicting senescence in ejaculate traits and positive values showing improvement in ejaculate traits with advancing male age. Bold error bars (C.I.) show whether overall effect size is significantly different from zero (*i.e.* not overlapping zero), while light error bars show the prediction interval (P.I.) of effect sizes. Sample sizes reported as: k = number of effect sizes (in brackets: number of studies).

### Aim 2: Role of biological and methodological moderators

#### Full model

We did not find a significant general effect of advancing male age on ejaculate traits (mean [95% confidence interval]: -0.197 [-1.496 to 1.103]), even when we included various moderators. These moderators were: taxonomic class, ejaculate trait, population type, proportion of maximum adult lifespan sampled, whether or not males had control over ejaculation, method of age estimation, sampling method, whether or not a study was experimental, and whether or not males underwent “unnatural” manipulations. However, the included moderators explained a significant proportion of the total heterogeneity our data (*R^2^* = 12.17%, QM = 99.606, QE_1765_ = 15299.075, P < 0.001, DF = 36).

### Role of individual biological moderators

#### Taxonomic class

We did not find evidence for age-dependent changes in ejaculate traits in any taxonomic classes (*R^2^* = 8.26%, QM = 26.082, P = 0.025, DF = 14, Fig. 1B for four major classes, Supplementary Fig. 3 for all classes), except in Malacostraca (which showed improvement with advancing male age).

#### Trait

We did not find any evidence for male age to affect individual ejaculate trait significantly when all animals in our dataset were considered (*R^2^* = 1.72%; QM = 51.287; P < 0.001, DF = 13, Fig. 2A). However, there were taxonomic class-specific (Trait x Taxonomic Class model: *R^2^*= 9.19%, QM = 130.65, P < 0.001, DF = 44, Fig. 3) and species-specific effects of male age on various ejaculate traits (Appendix 12). Specifically, for insects (Insecta, k = 258), ejaculate size (*i.e.* volume or weight of ejaculate), sperm concentration, quantity of sperm (corrected by body or testis size), number of sperm, and sperm viability improved with advancing male age (Fig. 2B). For fish (Actinopterygii, k = 174), sperm motility and velocity decreased, whilst ejaculate size increased with advancing male age (Supplementary Fig. 4A). In birds (Aves, k = 318), sperm from older males had higher levels of oxidative stress than sperm from younger males (Supplementary Fig. 4B). Finally, no significant net effects of advancing male age were found on any ejaculate traits for mammals (Mammalia, k = 990; Supplementary Fig. 4C).

**Figure 2:**
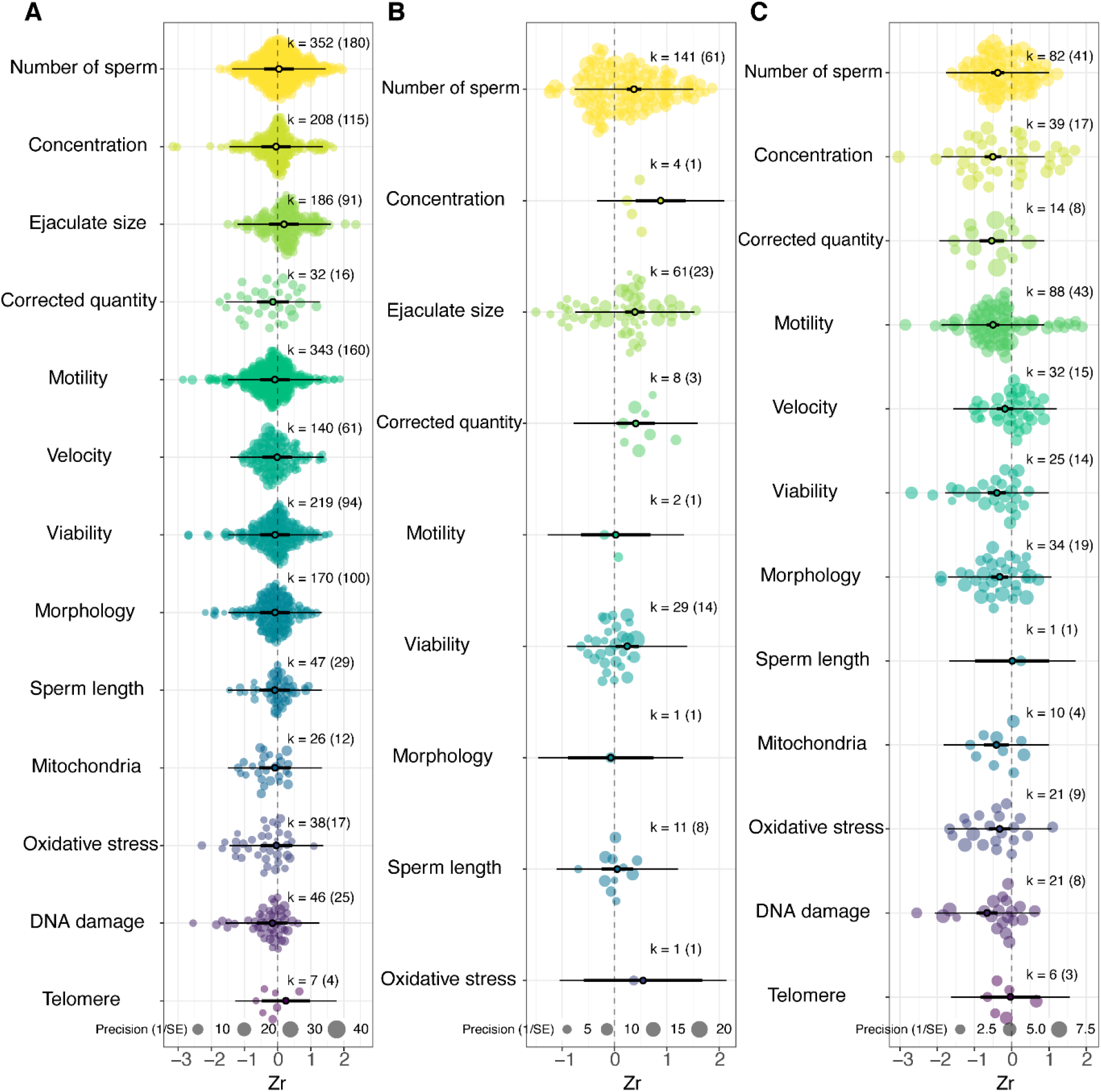
No consistent evidence for senescence in ejaculate traits when all taxa were considered, but some ejaculate traits showed senescence in lab rodents, yet others improve with age in insects. **A.** Effect of advancing male age on individual ejaculate traits across all 157 species in the dataset. **B.** Effect of advancing male age on individual ejaculate traits in the class Insecta. **C.** Effect of advancing male age on individual ejaculate traits for the two most over-represented species combined (lab rodents): *Mus musculus* and *Rattus norvegicus.* The size of each data point represents the precision of the effect size (1/SE). The X axis represents values of effect sizes as Fisher’s z-transformed correlation coefficient (Zr), while the Y axis shows the density distribution of effect sizes. The position of the overall effect is shown by the dark circle, with negative values depicting senescence in ejaculate traits and positive values showing improvement in ejaculate traits with advancing male age. Bold error bars (C.I.) show whether overall effect size is significantly different from zero (*i.e.* not overlapping zero), while light error bars show the prediction interval (P.I.) of effect sizes. Sample sizes reported as: k = number of effect sizes (in brackets: number of studies).

**Figure 3:**
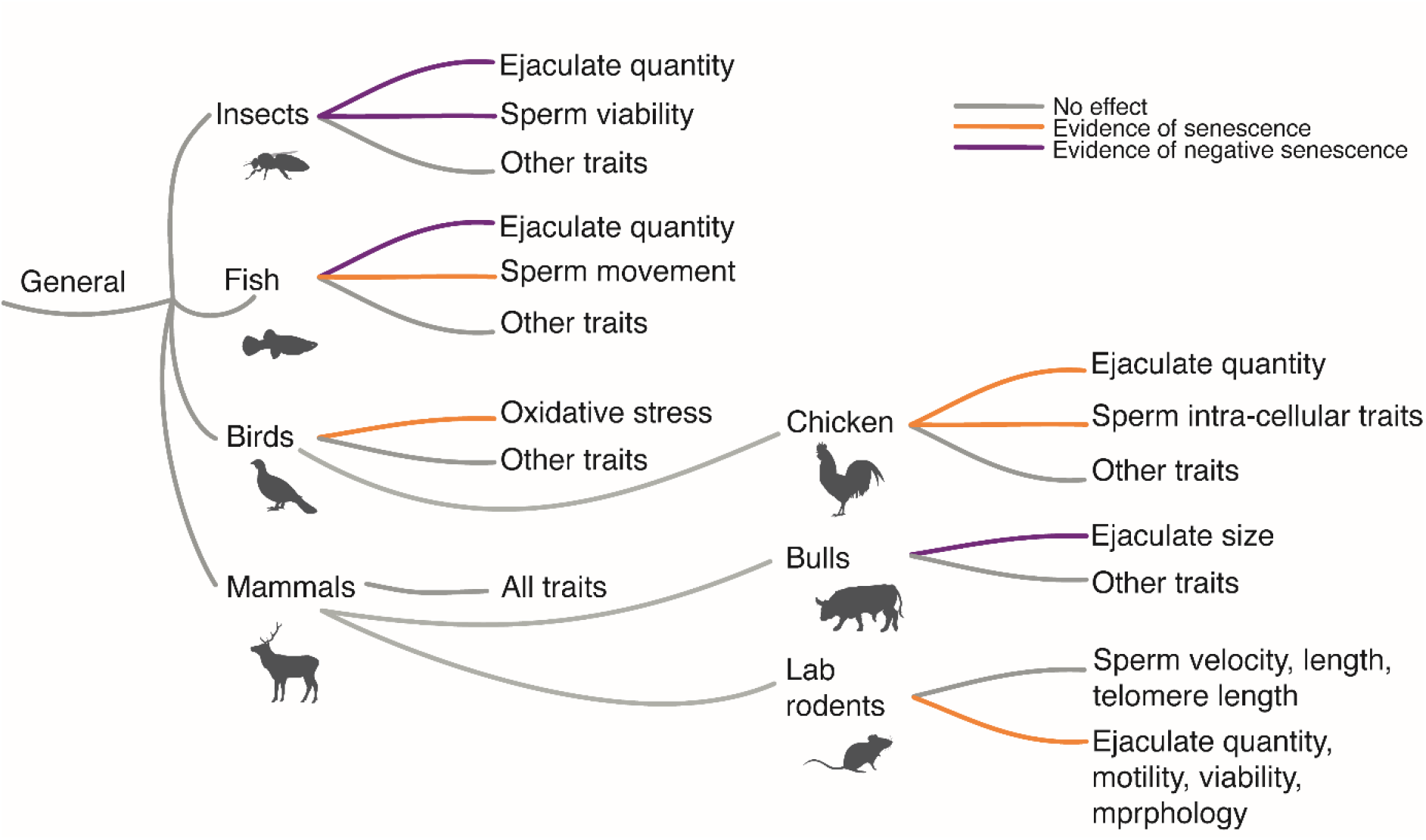
Specific ejaculate traits and taxonomy interacted to affect the evidence for senescence. Summary of results for how advancing male age affects different ejaculate traits across various taxa in our meta-analysis. Chicken refers to domestic chicken and red junglefowl.

We analysed four over-represented species in separate models (mice and rats, bulls, chicken), which in total represented ∼40% of the effect sizes in our dataset. For *Rattus norvegicus* and *Mus musculus* (k = 373, combined), most ejaculate traits (*i.e.* sperm viability, number, motility, morphological defects, concentration, mitochondrial function, DNA and oxidative damage) showed senescence with advancing male age (Fig. 2C). This evidence remained even when we only analysed males from studies that did not manipulate individuals and ‘controls’ that did not undergo “unnatural” manipulations (Supplementary Fig. 5A). For bulls (*Bos taurus,* k = 173), ejaculate size increased with advancing male age, but no other trait was affected significantly by male age (Supplementary Fig. 5B). For *Gallus spp.* (*i.e.* studies on domestic chicken and red junglefowl combined, k = 183), number of sperm, ejaculate size, sperm mitochondrial function, oxidative stress, and DNA damage, all showed senescence with advancing male age (Supplementary Fig. 5C).

#### Gonadosomatic Index (GSI)

The male GSI of a species (*i.e.* ratio of testes weight to body weight) did not influence how advancing male age affected ejaculate traits (*R^2^* = 0.26, QM = 0.786, P = 0.375, DF = 1, Supplementary Fig. 6).

### Role of individual methodological moderators

#### Manipulation

We compared the effects of male age on ejaculate traits in males that underwent unnatural manipulations (*i.e.*, experienced conditions outside of their typical range and had a control in the same study), to both, their ‘controls’ and males from studies that did not use unnatural manipulations (collectively called unmanipulated males). We found no overall evidence for senescence or improvement in ejaculate traits from either manipulated or unmanipulated males (*R^2^* = 0 %, QM = 0.021, P = 0.989, DF = 2, Supplementary Fig. 7A, B), or significant differences in effects sizes between manipulated and unmanipulated males (P = 0.885).

#### Population type

Population type did not modulate the effect of male age on ejaculate traits (*R^2^* = 1.12%; QM = 2.724, P = 0.605, DF = 4, Fig. 1C).

#### Sampling method

We found no overall evidence for senescence or improvement in ejaculate traits irrespective of whether studies sampled males longitudinally, semi-longitudinally, or cross-sectionally (*R^2^* = 0.08%, QM = 0.639, P = 0.887, DF = 3, Supplementary Fig. 8).

#### Ejaculate collection method

We did not find evidence for male age to affect ejaculate traits irrespective of whether males had control over ejaculation or not (*R^2^* = 1.36%; QM = 7.52, P = 0.023, DF = 2, Supplementary Fig. 9). However, males who had control over ejaculation exhibited a greater tendency for deterioration in ejaculate traits with advancing age, compared to males who did not have control over ejaculation (P = 0.006, DF = 1, Supplementary Fig. 9).

#### Proportion of maximum adult lifespan sampled

Studies sampling a higher proportion of maximum adult lifespan of the study organism provided stronger evidence for senescence in ejaculate traits across all animals (*R^2^* = 0.57%, QM = 4.838, P = 0.028, DF = 1, Fig. 4; see Supplementary Fig. 10 for distribution of LS sampled across taxa). When we analysed the four population types separately, we found evidence for senescence in ejaculate traits to increase with the proportion of lifespan sampled only in lab and captive populations (Fig. 4), but not in domestic or wild populations (Fig. 4).

**Figure 4:**
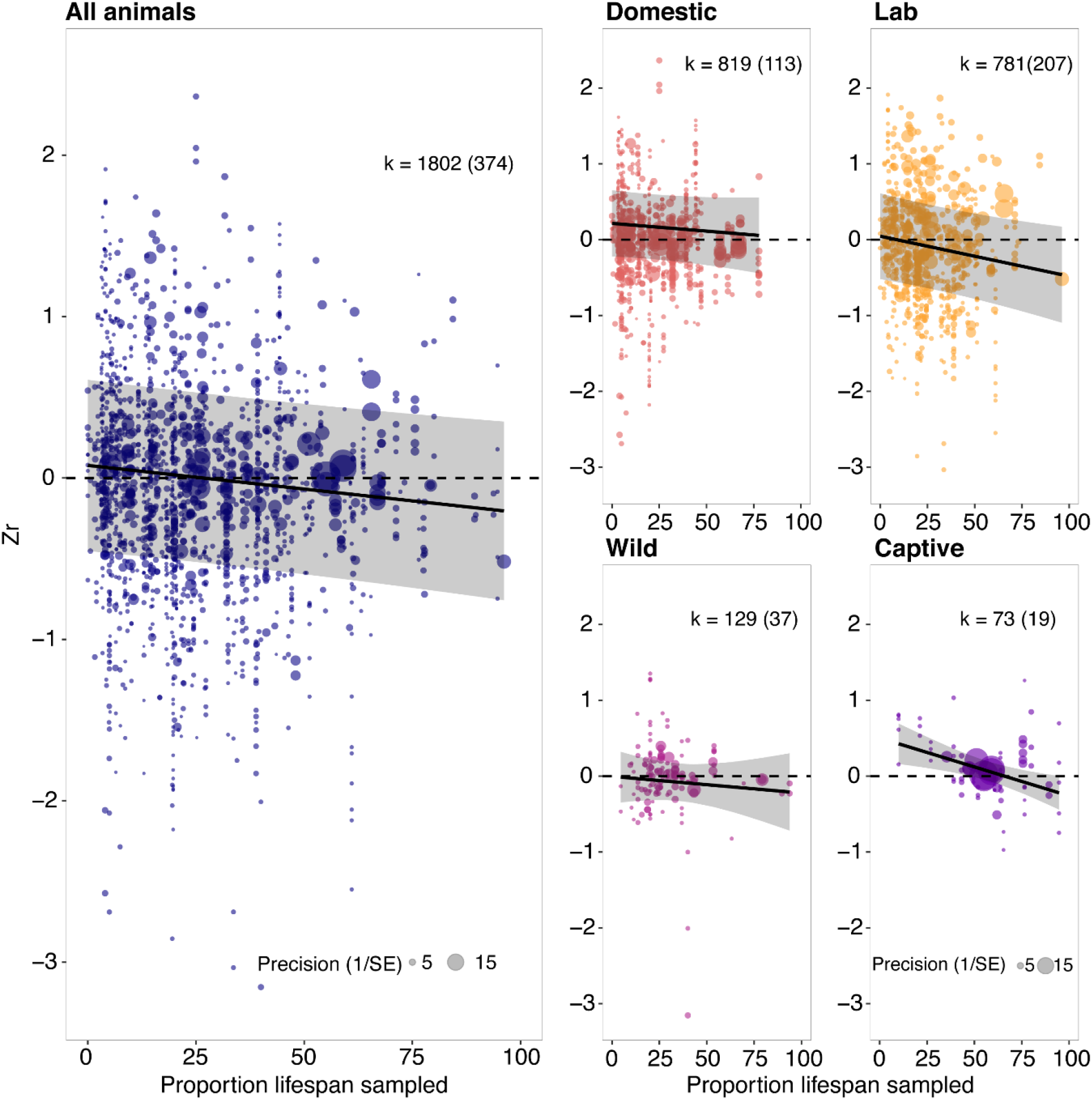
Increasing the proportion of maximum adult lifespan sampled, increased the evidence for senescence. Effect of proportion of maximum adult lifespan sampled (X axis) on the effect size *i.e.* Fisher’s z transformed r (Y axis) across all animals, and broken down for domestic, laboratory, wild, and captive animals. The size of each data point represents the precision of the effect size (1/SE). The dark line with shaded bars represents the overall effect of LS sampled on effect sizes and its 95% confidence intervals, respectively. Negative values depict senescence in ejaculate traits with advancing age while positive values show improvement in ejaculate traits with advancing male age. Sample sizes reported as: k = number of effect sizes (in brackets: number of studies).

#### Cold storage

We found no evidence for male age to affect ejaculate traits irrespective of whether or not ejaculates were stored in cold conditions (<5°C, regardless of the duration of storage) before analysis of sperm performance (*R^2^* = 0%, QM = 0.421, P = 0.810, DF = 2, Supplementary Fig. 11).

### Effects of advancing male age on reproductive traits (aim 3)

Male reproductive traits did not improve or decline with advancing male age (Supplementary Fig. 12A). However, male reproductive traits were less likely to deteriorate with advancing age than ejaculate traits (QM = 9.783, P = 0.002, DF = 1; Supplementary Fig. 12B).

### Other analyses

#### Quadratic effects of age

We found significant quadratic effects of [standardised] male age on [standardised] ejaculate traits (Supplementary Fig. 13). Specifically, we found significant quadratic effects for age-dependent changes in: percent morphologically normal sperm (t = -3.023, P = 0.003, DF = 346); percent motile sperm (t = 2.296, P = 0.022, DF = 589); and percent viable sperm (t = - 3.909, P < 0.001, DF = 437).

#### Publication bias

We found little evidence for publication bias. Visual inspection of funnel plots did not indicate any asymmetrical distribution of effect sizes around zero, indicating no evidence for publication bias (Supplementary Fig. 14, test for funnel plot asymmetry: P = 0.992). Our multi-level meta-regression publication bias test indicated no evidence for a small study bias (t = 7.21, P = 0.721, DF = 1810; Supplementary Fig. 15A) but significant evidence for a time-lag bias, with more recent studies being more likely to show reproductive senescence (t = -2.34, P = 0.019, DF = 1810, Supplementary Fig. 15B). We found no evidence for missing studies (missing studies on right= 0, SE = 10.94) in our trim-and-fill model that used one averaged effect size from each study (n = k = 379). Finally, we did not find a bias toward more significant effect sizes in our selection model (Supplementary Fig. 16).

## Discussion

Senescence lies at the core of life-history theory (Kirkwood and Rose, 1991; Stearns, 1989). Thus, we cannot fully understand organismal biology, and evolutionary processes such as sexual selection and sexual conflict (Archer and Hunt, 2015; Bonduriansky et al, 2008; Graves, 2007; Maklakov et al, 2009), without understanding the causes and consequences of male reproductive senescence (e.g. Archer et al, 2018). Here, we investigated overall patterns of age-dependent changes in ejaculate traits across [non-human] male animals, how these patterns are modulated by various biological and methodological factors, and how changes in ejaculate traits with advancing male age compare against changes in reproductive traits. Overall, we found no general pattern of senescence, however we suggest that this is likely due to methodlogical limitations in studies, and biological differences between taxa.

### Aim 1: Effects of advancing male age on ejaculate traits

Contrary to expectations, we found no consistent general pattern of senescence in ejaculate traits across studies. We suggest several potential non-mutually exclusive reasons for this. First, while we found that increasing the proportion of lifespan sampled by a study yielded greater evidence for senescence (also shown by Fricke et al, 2023; Vrtilek et al, 2022), studies tended to sample a low proportion of maximum adult lifespan (median = ∼25%, Supplementary Fig. 10). Limiting sampling to early-life of organisms may have led to an underestimation of senescence in our meta-analysis, as senescence is predicted to be observed later in life (Gaillard and Lemaitre, 2020; Fricke et al, in prep., Jones et al, 2014; Kirkwood and Rose, 1991; Lemaitre et al, 2020; Peron et al, 2010; Williams et al, 2006). One reason for such limited sampling of older males could be associated with many studies not explicitly testing for senescence. Instead, some of the studies in our dataset were interested in estimating peak reproductive ages of males (e.g. Hallap et al, 2006), improving age-related reproduction (e.g. Hocking and Bernard, 1997), testing the effects of pharmacological interventions (e.g. Marcinak et al, 2017; Zhao et al, 2020), or comparing methods for sperm preservation (e.g. Garcia et al, 2017). We suggest future studies to explicitly test for senescence and sample males to later life to help elucidate patterns of senescence more accurately.

Second, age-dependent changes may not be linear (Baudisch 2011; Baudisch et al, 2013; Jones et al, 2014), but rather follow a curvilinear (e.g. convex) pattern. Our test of quadratic effects showed some evidence in support of this. Thus, if ejaculate traits improve from early to mid-adult life (e.g. Cooper et al, 2021) and deteriorate (*i.e.* senesce) only later in life, the positive part of the function would be disproportionately weighted against the negative part of the function (Pizzari et al, 2008). This is because there are usually fewer older males sampled than younger males, as evidenced by the focus of studies being mostly restricted to 25% of maximum organismal lifespan. Curvilinear patterns of age-dependent changes in ejaculate traits would lead most studies to underestimate senescence and result in an overall positive or net zero effect, when age-dependent changes are analysed as a linear function (e.g. using effect sizes). We suggest that studies on age-dependent changes in reproductive traits analyse and report the shape, rate, and onset of senescence (see Jones et al, 2014) whenever possible, and that meta-analysts introduce techniques where non-linear relationships can be analysed.

Third, most studies in our dataset were conducted cross-sectionally, where different cohorts of males were assigned to different age groups, and measured once. Cross-sectional sampling can underestimate senescence when there is selective disappearance of poor-quality individuals with increasing age (Hamalainen et al, 2014; Sanghvi et al, 2022). Cross-sectional sampling could lead to a biased sampling of good quality males in older age cohorts. We did not find significant evidence for senescence in ejaculate traits within longitudinal studies either. However, to account for selective disappearance in longitudinal studies properly, one would need to analyse individual-level data for each male in each study rather than the means of age groups, and these data are seldom reported (Nussey et al, 2011; van de Pol and Verhulst, 2006).

Fourth, terminal investment by old males may also contribute to masking the effects of ejaculate deterioration, such that older males, who have lower survival and reproduction prospects in the future, invest more in their current reproduction than young males (Clutton-Brock, 1984). Age-dependent terminal investment has been shown in only a few taxa (e.g. Duffield et al, 2018; Froy et al, 2013; Part et al, 1992; Tarwater and Arcese, 2017), and whether it is common across animals is yet unknown.

Fifth, negligible senescence or age-dependent improvement in ejaculate traits in our study could also be due to reproductive senescence not being a general outcome of ageing (Finch, 2009; Jones et al, 2014; Jones and Vaupel, 2017; Roper et al, 2021; Shefferson et al, 2017), and some animals showing true negligible (also suggested for plants: Munne-Bosch, 2015) or negative senescence (Vaupel et al, 2004). The high heterogeneity (𝐼^2^= 95%) in study outcomes seen in our meta-analysis indicates that studies some show both true senescence or negative senescence in ejaculate traits.

### Aim 2: Effects of individual moderators

Some biological and methodological moderators were important in modulating the effects of advancing male age on ejaculate traits. As the effects of these moderators were tested individually, our results could possibly be explained by other confounding factors or by moderators not simultaneously included in our meta-analysis. These results should thus only be treated as hypothesis generating rather than causal.

#### Biological moderators

Taxonomic class was an important modulator of the effects of advancing male age on ejaculate traits. In comparative studies, taxonomy has been shown to play a key role in explaining differences in senescence patterns between different animal groups (Jones et al, 2008; Jones et al, 2014; Reinke et al, 2022). This can be attributed to different ecologies, niches, behaviours, life-history strategies, metabolisms, and evolutionary histories of animals. For instance, vertebrates often show higher levels of paternal care than invertebrates (Balshine, 2012; Clutton-Brock, 1991; Reynolds et al, 2002; Royle et al, 2012; Suzuki, 2013; Trumbo, 2012). Thus, male vertebrates may have evolved to invest more in somatic maintenance and provisioning, and less in ejaculate traits, than invertebrate males. Additionally, taxa that have continuous growth (e.g. fish), may invest more in somatic maintenance, than taxa with determinate growth (Patnaik et al, 1994). This could lead to greater evidence for reproductive senescence, but lower actuarial senescence, in taxa that invest more in growth and somatic maintenance than other taxa (Hoekstra et al, 2020; Vrtilek et al, 2022).

For some taxa, we found that effects of advancing male age differed between various ejaculate traits (Fig. 3). Patterns of reproductive senescence have previously been shown to differ between reproductive traits within a given clade (Hayward et al, 2015; Naciri et al, 2022). This could be due to: covariances between different traits (Snook, 2004); some traits being more sensitive to age-dependent deterioration than others (Cooper et al, 2021; Sanghvi et al, 2021); or traits being under different selection pressures (Moorad and Ravindran, 2022; Schluter et al, 1991). For example, studies have shown that sperm movement negatively correlates with sperm longevity (Cardozo et al, 2020; Levitan, 2000; Smith, 2012) and positively with oxidative stress (Fasel et al, 2017). Similarly, sperm quality can trade-off with sperm quantity (Evans, 2011; Liao et al, 2019; Moore et al, 2004; Snook, 2005), although see Gomez-Montoto et al (2011) for contrasting evidence. Additionally, some traits are likely to be more important than others for certain species in driving fertilisation success (Birkhead et al, 1999; Gage and Morrow, 2003; Gage et al, 2004; Hunter and Birkhead, 2002; LaMunyon and Ward, 1998; Radwan, 1996; Simmons, 2003; Snook, 2004; reviewed in Simmons and Fitzpatrick, 2012). For instance, Gage and Morrow (2003) show that increased sperm number, but not sperm size, determines fertilisation success in crickets. On the other hand, increase in sperm velocity, but not number, longevity, or length, leads to higher fertilisation success in salmon (Gage et al, 2004). As a consequence, traits that are more important determinants of male fertilisation success for specific taxa could be selected to deteriorate at lower rates with age, at the cost of higher deterioration in less-important determinants of fertilisation success (Moorad and Ravindran, 2022).

Insects in our study showed a significant increase in all sperm quantity traits, *i.e.* sperm number, concentration, and ejaculate size. This increase could be associated with their mating status, as most studies (>75%) on insects in our meta-analysis kept males as virgin (see Fricke et al, 2023; Richardson and Zuk 2022, who show similar results in males and females, respectively). Low mating rates could result in old insects accumulating more sperm and producing larger ejaculates than young males, because many insects have continuous spermatogenesis throughout their lives (Bressac et al, 2009; Hiroyoshi et al, 2021; Malawey et al, 2019; Reinhardt et al, 2011). In support, Fox et al (1995), Kehl et al (2013), and Sepil et al (2020) show that age-dependent decreases in insect reproductive output can be masked by accumulation of ejaculates in old virgin males. Another explanation could be that male insects reach sexual maturity many days after eclosion (e.g. Pitnick 1993), thus sampling early- to mid-life would only reflect the maturation period, and consequently improvement, in insect ejaculates. While improvement in ejaculate quantity traits could be partly explained by mating status or lifespan sampled, they do not help explain our finding on improvement in sperm viability, which future work could address.

Fish (Actinopterygii) showed some evidence for senescence in sperm performance traits (*i.e.* velocity and motility), but also age-dependent increases in ejaculate size. Some studies on fish have shown that older males have more sperm due to age-dependent accumulation of sperm when males are seldom mated (e.g. Aich et al, 2021; Vega-Trejo et al, 2019). To test this hypothesis in fish, we suggest that future studies should include information on male mating history to enable a more informed analysis of reproductive senescence. Another possible explanation is that old fish could be investing in producing larger ejaculates to compensate for senescence in sperm performance traits. Future studies could test whether covariances between sperm performance and ejaculate size in fish are modulated by male age. Age-dependent improvement in ejaculate size could also be due to continuous post-maturity growth in fish body size (Zak and Reichard, 2020; Vrtilek et al, 2022), leading to older males having larger gonads and being able to produce larger ejaculate volumes than young males. Lastly, age-dependent increases in ejaculate size could be due to an increase in seminal fluid production with age, rather than an increase in sperm number, which could be investigated by future studies. Our results are in line with a recent review on fish (Vrtilek et al, 2022), which shows evidence for male reproductive senescence in less than half of the studies analysed.

We did not find consistent evidence for senescence in ejaculate traits across mammals. However, we found taxon-specific senescence or age-dependent improvement in ejaculate traits of some mammalian species. In lab rodents (*i.e.* mice (*Mus musculus*) and rats (*Rattus norvegicus*) combined), we found significant evidence for senescence across a wide range of ejaculate traits (even when only control/wild type genetic strains were analysed, e.g. WT and C57 for mice, Brown Norway and Sprague Dawley for rats). The evidence for consistent senescence in ejaculate traits could be partially attributed to these species being kept in highly controlled environments. For bulls, we found an increase in ejaculate size with age. This could be due to studies on bulls being conducted on commercial farms, where older males are usually kept solely for breeding purposes. Here, older bulls with low ejaculate production are culled (Khatun et al, 2013; Wondatir-Workie et al, 2021), thus studies on bulls could be representing highly biased cohorts of old males.

We did not find evidence for senescence in ejaculate traits in birds, except for age-dependent increase in oxidative stress to sperm. Yet, in studies of domestic chicken and red junglefowl combined (*Gallus spp.*), which represented >50% of bird effect sizes, we found evidence for senescence in ejaculate quantity traits (*i.e.* sperm number, ejaculate size), and in various intra-cellular aspects of sperm. Despite studies on *Gallus spp.* sampling on average a low proportion of maximum adult lifespan (∼10% median), senescence in ejaculate traits was detected. This might indicate early onset of senescence due to domestication for fast life-histories (also suggested by Bergeron et al, 2023), because domestic chickens, especially breeds like broilers, have been artificially selected for extremely short generation times (Bennett et al, 2018; Hartcher and Lum, 2020).

#### Methodological moderators

Extending the proportion of maximum adult lifespan sampled, increased the evidence for senescence in ejaculate traits, of a species. This result suggests that the onset of reproductive senescence generally occurs late in life (Charlesworth, 1993; Gaillard and Lemaitre, 2020; Fricke et al, 2023; Jones et al, 2014; Lemaitre et al, 2020; Monaghan and Metcalfe, 2019; Peron et al, 2010). Thus, senescence will more likely be detected if studies sampled a larger proportion of lifespan.

Ejaculate collection method explained significant heterogeneity. Males that had control over ejaculation showed more age-dependent deterioration in ejaculate traits than males that did not have control over ejaculation (although neither showed overall improvement or senescence with advancing male age). One possible explanation is that in species characterised by temporal sperm stratification in the male extragonadal sperm reserves (e.g. vas deferens), control over ejaculation may enable males to utilise sperm largely from the most distal part of the sperm reserves, which are farthest away from the testes. These sperm would be older and thus could have incurred higher post-meiotic damage (Pizzari et al, 2008; Reinhardt, 2007). On the other hand, studies that dissect, electro- ejaculate, massage, or “strip” males, may obtain sperm from the entire reproductive duct, including younger sperm. These invasive methodologies may therefore yeild ejaculates with lower net post-meiotic damage, although also with immature sperm (in species with extra- gonadal sperm maturation). Future studies can test these hypotheses by comparing sperm across different sections of the male extra-gonadal sperm reserves, and testing if male sampling method can causally affect the evidence for senescence in ejaculate traits (e.g. Axner et al, 1997; reviewed in Reinhardt, 2007).

Contrary to our predictions, population type did not modulate patterns of senescence in ejaculate traits. Each population type could influence male reproductive senescence in contradicting ways (Reichard, 2016; Ricklefs, 2008), depending on interactions with other moderators. For instance, wild populations usually live in environments with greater food and climate uncertainty, competition, predation, disease, and physical activity, thus can be expected to show more reproductive senescence than captive animals (Cooper and Kruuk, 2018; Nussey et al, 2013; Tidiere et al, 2016; Soulsbury and Halsey, 2018). On the other hand, individuals in wild populations usually face higher extrinsic mortality and may not survive long enough to be sampled at older ages when senescence is manifested (Hamalainen et al, 2014; Xia and Moller, 2022). Accurately testing for effects of population type would require building more nuanced models that consider such interactions, something we were unable to test due to lack of such data being consistently reported across population types.

### Aim 3: Effects on male reproductive traits

We found no consistent evidence for overall improvement or senescence in reproductive traits of males (*i.e.* measures of fertilisation success, egg/offspring number/viability/quality). However, reproductive traits were less likely to exhibit age-dependent deterioration than ejaculate traits, for which there could be several reasons. First, not all ejaculate traits may be important in determining reproductive output. Thus, deterioration in some ejaculate traits might have little consequence for reproductive traits such as fertilisation success (Gage et al, 2004). Second, if reproductive output is condition-dependent (Kokko, 1998) with positive covariances between survival and reproduction, old males might have higher reproductive output due to viability selection (Brooks, 2001; Johnson and Gemmell, 2012). Third, female-driven effects (e.g. cryptic female choice, plastic maternal investment in eggs) could buffer male reproductive senescence (Firman et al, 2017; Harris and Uller, 2009). Fourth, evidence for senescence in male reproductive traits may manifest in offspring quality (for which we had limited data) rather than quantity. Fifth, senescence in pre-copulatory male traits might affect male reproductive traits more than ejaculate traits do (Gasparini et al, 2019). Lastly, our meta-analysis used data on reproductive traits only from studies that also measured ejaculate traits, which may be a small and biased representation of data.

Senescence in male reproductive traits can have serious consequences for female fitness, influence sexual selection on males (Archer and Hosken, 2023; Promislow, 2003), and lead to sexual conflict (Adler and Bonduriansky, 2014; Bonduriansky et al, 2008; Dean et al, 2010; Radwan, 2003). However, age-dependent changes in ejaculate traits may not always accurately reflect changes in reproductive traits. Thus, it is essential that future studies on male senescence measure ejaculate, reproductive, and offspring traits, to get an unbiased view of the fitness consequences of male ageing.

## Conclusion

Our meta-analysis aimed to understand broad patterns of senescence in ejaculate traits. We did not find consistent evidence for male reproductive senescence in animals (*aim 1*). This is likely due to biological differences and various methodological limitations across studies, rather than reflect a genuine absence of senescence. For instance, studies rarely accounted for male mating history, and usually sampled low proportions of an organism’s lifespan.

Curvilinear shapes of ageing and selective disappearance of males could also underestimate the evidence for reproductive senescence. We also investigated how different moderators affect age-dependent changes in ejaculate traits (*aim 2*) and found that reproductive senescence may be trait (e.g. Naciri et al, 2022) and taxon-specific (e.g. Jones et al, 2014). Lastly, we found some evidence for advancing male age being less detrimental to male reproductive traits than to ejaculate traits (*aim 3*).

Our meta-analysis includes data from species that are model organisms in medicine and gerontology (e.g. lab rodents), animal husbandry and theriogenology (e.g. bulls and chicken), evolutionary ecology (insects, birds, wild and lab animals), conservation (animals in zoos and captive breeding programs), and aquaculture (fish, crustacea). In addition to contributing to our understanding of reproductive senescence, the results of our study have therefore relevance across a variety of fields, and may help informing breeding programs for fisheries, commercial farms, and zoos, and perhaps help us understand human reproductive senescence better. Overall, our study identifies methological improvements and suggests novel hypotheses for studying senescence.

## Acknowledgements

We are extremely grateful to all the researchers who provided us with unpublished/missing data and life history tables for various species, without whom our meta-analysis would not be possible: Abdallah Assiri, Adolfo Cordero, Adrienne Crosier, Alberto Velando, Alfonso Bolarin, Aline Malawey, Alistair Senior, Anders Pape Moller, Anil Kumar, Antje Girndt, Ashley Watt, Asim Orem, Bradley Metz, Bryan Neff, Budhan Pukazhenthi, Charles Fox, Chris Friesen, Chris Weldon, Christine Miller, Christophe Bressac, Claudia Fricke, Claudio Maia, Claus Wedekind, Clelia Gasparini, Clint McDonald, Craig Packer, Daniel Sasson, Davnah Payne, Diana Perez Staples Folger, Martha Reyez Hernandez, Distl Ottmar, Elena Zambrano, Emily “Becky” Cramer, Emily Duval, Erin MacCartney, Felipe Martinez, Fumio Hayashi, Gabriele Sorci, Gerard Wilkinson, Gerlind Lehmann, Gregor Majdic, Hasan Sevgili, Heriberto Martinez, Ilie Racotta, Ioannis Tsakmakidis, Jan Lifjeld, Jane Hurst, Janice Bailey, Maurice Clotilde, Jesus Dorado, Jurgen Heinze, Karen Lockyear, Karolina Stasiak, Katarzyna Kotarska, Kathrin Langen, Klaus Reinhardt, Leandro Miranda, Leslie Curren, Linda Penfold, Maira Brito, Malgorzata Kruczek, Manasi Kanuga, Mark Elgar, Martin Brinkworth, Martin Schulze, Maud Bonato, Megan Head, Melissah Rowe, Michael Greenfield, Michael Ritchie, Michele Di Iorio, Michelle Helinski, Milos Krist, Moira O’Bryan, Muhammed Ines Inanc, Naomi Pierce, Nicolaia Iaffaldano, Nikos Papadopolous, Nikron Thongtip, Nils Cordes, Nucharin Songsasen, Megan Brown, Pablo Bermejo Alvares, Paco Garcia Gonzales, Panos Milonas, Patricia Diogo, Paul Joseph, Philip Downing, Priscilla Ramos, Rakesh Seth, Stuart Reynolds, Rebecca Dean, Sachiko Koyama, Satoshi Hiroyoshi, Silvia Cerolini, Sina Metzler, Stanislaw Kondrack, Stefan Luepold, Steven Ramm, Stuart Meyers, Theo Bakker, Marion Mehlis, Tobias Kehl, Triin Hallap, Ulrike Luderer, Upama Aich, Wael Farag, Wei Shi, Wen Liao, Xiaoxu Li, Yasaman Alavi, Yih Fwu Lin, and Yingmei Zhang.

We are also thankful to Milan Vrtilek, Rose O’Dea, Alex Kacelnik, Ana Silva, and Ellie Bath for their helpful comments and suggestions.

## Author contributions

KS, TP, IS conceptualised and designed the study. KS, SG, RVT screened the studies. SN, SJ provided extra studies. RSG suggested methods to standardise age. KS extracted data. RVT checked for repeatability of data extraction. RVT, SN, SJ, KS wrote the code and analysed the data. KS, RVT, IS, TP wrote the first draft of the manuscript. All authors contributed in critical assessment of the manuscript and subsequent revisions.

