## Appendix for "No general effects of advancing male age on ejaculates: a meta-analysis across the animal kingdom"

**Appendix 1:** Words used to address the question of how advancing male age affects ejaculates in non-human animals.

The table shows the most frequent words from our scoping search collected from 48 relevant abstracts generated through a word cloud. Words used in our search string are highlighted in bold and italics. The selection of words was made such that they were not specific to a single species, nor biologically broad.

| frequency | word | frequency | word |
|---|---|---|---|
| **305** | ***sperm*** | 25 | size |
| **191** | ***male*** | 24 | fluid |
| **151** | ***age*** | 24 | transfer |
| 104 | mating | 22 | fitness |
| 75 | reproductive | 21 | offspring |
| 66 | female | 21 | proteins |
| 60 | females | 20 | competitive |
| 58 | success | 20 | ejaculates |
| 53 | competition | 20 | fertilization |
| **47** | ***ejaculate*** | 20 | mated |
| **45** | ***old*** | 20 | selection |
| **45** | ***young*** | 20 | transferred |
| 41 | quality | 19 | history |
| 36 | traits | 19 | Sperm |
| 33 | older | 18 | viability |
| 32 | seminal | 17 | associated |
| 31 | species | 17 | rate |
| 30 | ***ageing*** | 16 | Drosophila |
| 29 | fertility | 16 | paternity |
| 29 | sexual | 16 | production |
| **28** | ***senescence*** | **16** | ***semen*** |
| 27 | mate | **14** | ***aging*** |
| 25 | ability | 14 | cells |

**Appendix 2:** Search string

Distinct Boolean characters used for the search strings in each database.

Search string used for **SCOPUS**:

“( TITLE-ABS-KEY ( *male* ) AND TITLE-ABS-KEY ( *age* ) OR TITLE-ABS-KEY ( *ageing* ) OR TITLE-ABS-KEY ( *aging* ) OR TITLE-ABS-KEY ( *senesc** ) OR TITLE-ABS-KEY ( *old* ) OR TITLE-ABS-KEY ( *young* ) AND TITLE-ABS-KEY ( *sperm* ) OR TITLE-ABS-KEY ( *ejaculate* ) OR TITLE-ABS-KEY ( *semen* ) AND NOT TITLE-ABS-KEY ( *human* ) OR TITLE-ABS-KEY ( *men* ) ) “

Search string used for **Web of Science**:

“AB= (male AND (age OR ageing OR senesc* OR aging OR old OR young ) AND ( sperm OR ejaculate OR semen ) NOT ( human OR men) ) OR TI= (male AND (age OR ageing OR senesc* OR aging OR old OR young ) AND ( sperm OR ejaculate OR semen ) NOT ( human OR men) )”

Search string used for **BASE**:

“male age sperm”

**Appendix 3:** PRISMA flow diagram of data screening, identification, and inclusion procedure.

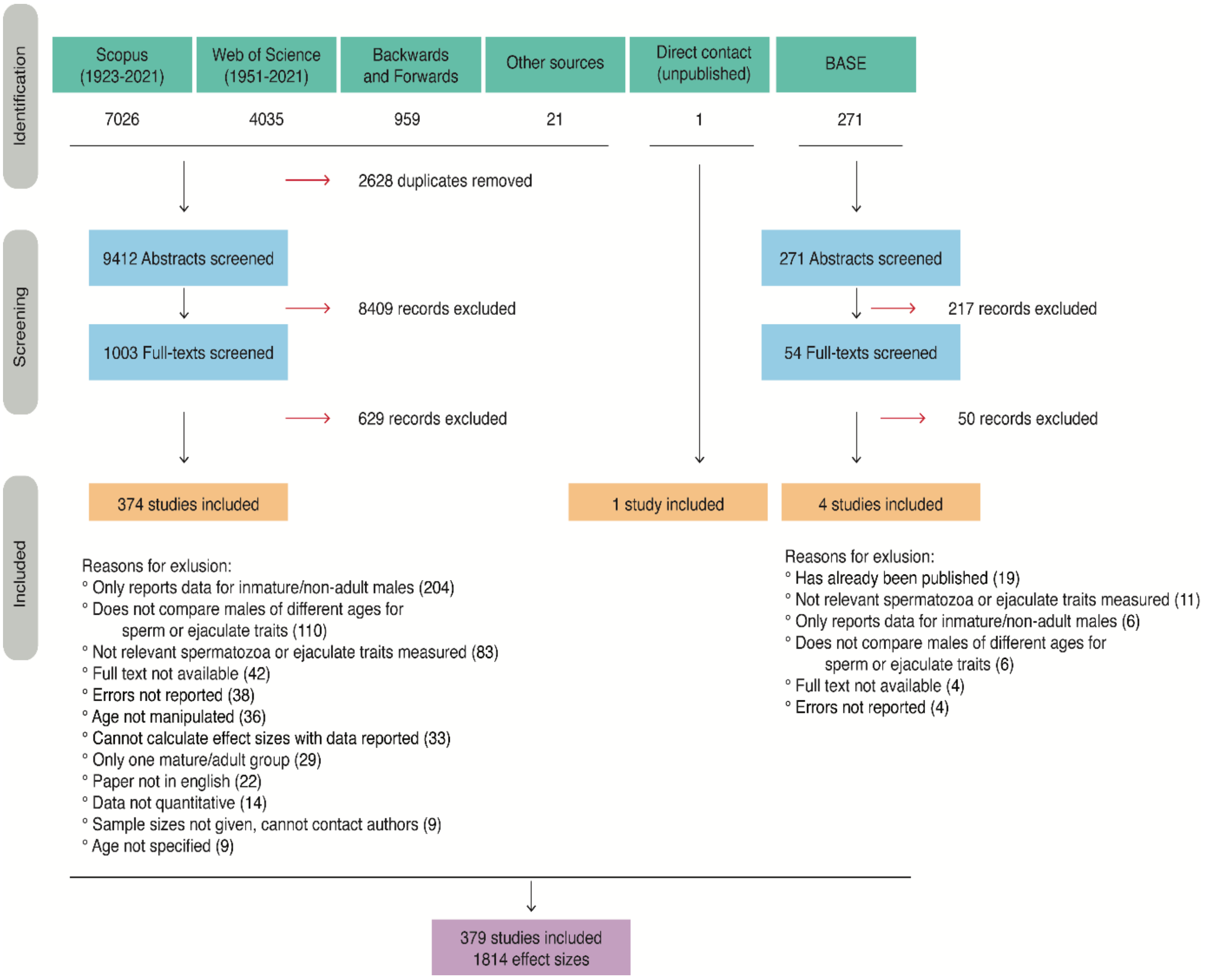

**Appendix 4:** Definitions of population type (i.e. setting)

Captive: Managed study population which lives in an enclosed environment, and used in a captive breeding program, captive breeding centre, wildlife conservation research centre, or zoo. The webpage of the reported institutes where the study was conducted was checked when unsure, to ascertain whether it fell into this category.

Lab: Study population maintained under or adapted to lab conditions (for at least their adult life), living in an enclosed environment (even if collected in the wild pre-adulthood), but not used for a captive breeding program.

Domestic: Study population descended from a line of deliberately artificially selected individuals (inferred), or raised on commercial farms whose purpose is using animals for direct human consumption (e.g. as food, clothing, protection).

Wild: Was born in the wild/lives in natural unenclosed environments and was caught from the wild (post-adulthood) for the study.

**Appendix 5:** Collection of data on adult lifespan, age of sexual maturity, and sperm competition levels for different species

If the study did not report the maximum adult lifespan or age of sexual maturity/adulthood of the species/population being studied, these were collected from other sources, and then used to calculate the percentage of maximum adult lifespan sampled for a given species in the study. Data on maximum adult lifespan of a given species, as well as age of sexual maturity were then used to calculate the proportion of maximum adult lifespan sampled for a given species in a study as follows:

$$Proportion\ of\ maximum\ adult\ lifespan\ sampled = \frac{\big((Max\ age\ sampled - \ age\ of\ maturity) - \ (Min\ age\ sampled\ - \ age\ of\ maturity)\big)}{(Maximum\ lifespan\ of\ species\ - \ age\ of\ adulthood)} * \ 100\%$$

To collect data on maximum lifespans (MAL) and age of sexual maturity/adulthood (AoSM) on a given species or population, we first prioritized using the data on lifespan reported directly in the study. If the study did not report MAL or AoSM, we collected this data from multiple sources. These included large publicly available databases such as Animal Diversity Web (https://animaldiversity.org/), AnAge (De Maghalhaes and Costa, 2009), Pantheria (Jones et al, 2009), and COMADRE (Salguero-Gomez et al, 2016), as well as from data in published articles that contained supplementary datasets on MAL and AoSM estimates of species included in our study. There were very few databases that contained MAL and AoSM for animals such as insects. As a result, peer reviewed, published articles in scientific journals were used for data on MAL and AoSM for such species. When values were reported as a survival curve, the maximum age of the curve where individuals were alive was used. When data was not available from any of these sources, we also contacted the corresponding authors of the papers in our meta-analysis for maximum lifespan estimates on the species. We only collected data for MAL and AoSM on males. When male-specific data was not available, species-specific data was collected. If a species had data on MAL and AoSM from multiple sources, these estimates were averaged, and the average was then used as the maximum lifespan of the species, so that exceptionally large values, which might be unusual for a species (e.g. red junglefowl maximum lifespan is reported as 30 years on AnAge, which is quite unusual), did not bias our dataset. Additionally, when data from multiple sources was available, we ensured that we used data from males which most closely matched the rearing conditions and morphs of the species/study males in our meta-analysis. For insects, age of adulthood was considered as the age of eclosion/ last moult. Note that for species which had domesticated as well as wild-derived populations in our dataset (e.g. red junglefowl vs chicken, both of which were called *Gallus gallus* in our phylogeny), we collected their lifespan and sexual maturity data separately.

We tested the association between maximum and average lifespans collected for each species. This was high (R sq. = 0.85). Thus, maximum lifespan was used because it was available for a greater number of species than average lifespan, and less affected by juvenile mortality.

We also recorded the level of sperm competition faced by a species by collecting data on the Gonadosomatic index (i.e. testis weight as a percentage of body weight), which has been shown to be a reliable predictor of sperm competition. This was done by searching for “testes weight” or “gonadosomatic index” of each species in our dataset on google scholar as well as collecting data from comparative studies on this topic (OSF https://osf.io/dk8sq/ for raw data).

**Appendix 6:** Trait description

For all studies, we collected data on ejaculate traits as reported by the original paper. When a study reported data for multiple traits, data on all traits were collected. Although, when a study reported data on a whole trait (e.g. % of total motile sperm; % sperm with morphological defects) as well as sub traits (e.g. % progressively motile sperm, % sperm with mid-piece defects), only data from the whole trait was recorded. Due to differences in terminology when describing similar traits between different studies, we created a broad category to describe different types of traits. The categorization of these traits is described below.

Ejaculate traits

| **In meta-analysis** | **In paper** |
|---|---|
| Concentration | Sperm density, concentration, count per mL/ µL, spermatocrit |
| DNA damage | Sperm chromatin damage, sperm DNA damage, sperm chromatin instability, sperm DNA fragmentation, sperm chromatin structure damage |
| Ejaculate size | Spermatophore size, ejaculate mass, ejaculate volume, area of seminal vesicle filled with ejaculate |
| Corrected quantity | Sperm concentration or number as proportion of body mass/testis mass/epididymis mass |
| Mitochondria | Sperm ATP, sperm metabolic activity, sperm mitochondrial function, sperm mitochondrial activity, sperm mitochondrial membrane potential |
| Morphology | Normal/abnormal sperm morphology, tail/head/midpiece morphological defects, cytoplasmic droplets on sperm |
| Motility | Percent progressive motility, percent non-progressive motility, percent total motility, sperm vigor, sperm mass motility, motility (on a numerical subjective scale) |
| Number of sperm | Number of apyrene sperm, number of eupyrene sperm, number/count of sperm, number of spermatophores, number of sperm bundles |
| Oxidative stress | Reactive oxygen species in sperm, sperm oxidant or antioxidant levels, sperm lipid peroxidation, sperm glutathione peroxidase, sperm superoxidase dismutase |
| Sperm length | Total sperm length, sperm tail length |
| Telomere | Sperm telomere length |
| Viability | Sperm acrosome integrity, sperm viability, % sperm alive/dead, sperm vitality, sperm membrane integrity |
| Velocity | Average path velocity (VAP), curvilinear velocity (VCL), straight line velocity (VSL) (in that order of preference, when more than one reported) |

Reproductive output traits

| In meta-analysis | In paper |
|---|---|
| Fertilization success | Percent eggs fertilized, Percent eggs sired under sperm competition, fertilization rate, fertilization success, PVL hole number, paternity share |
| Number/viability/quality of egg/offspring | Egg hatchability, egg mass, egg viability, male or female fecundity, hatching success, number of eggs laid, female lifetime reproductive success, number of offspring, number of fetuses, offspring developmental rate, offspring survival, offspring body mass/ size |

**Appendix 7:** “Unnatural” manipulations

“Unnatural” manipulations were defined as conditions experienced by males outside their physiological range (as defined in the study) and as conditions not typically experienced by healthy individuals (in the study populations). These manipulations also had a well-defined control in the study. These are as follows: pharmacological interventions such as toxins, chemicals, or medicines (Control: no pharmacological intervention); temperature manipulations (Control: standard temperature as defined in the study); radiation (Control: no radiation); genetic mutations/mutant lines with specific knocked out genes (Control: Wild type, or control lines as defined in the study); 24 hour dark or 24 hour light circadian durations (Control: 12:12 hour light durations); infection/disease (Control: no infection or disease); inbreeding (Control: Outbreeding); dietary or protein restriction (Control: Standard diet as defined in the study).

Other types of manipulations, such as sperm storage durations, mating history of males, seasons, male social status, or female age were not considered to be outside the typical range of conditions experienced by males, nor did they have an easy to define control, thus were not defined as “unnatural”.

**Appendix 8**: Moderators tested in our meta-regression models and their ranges/levels. For reasons for why these moderators were chosen, and how we expect them to affect male reproductive ageing, see Table 1 in main text.

| Moderator | Levels |
|---|---|
| Proportion maximum adult lifespan sampled | 0 to 100% |
| Ejaculate collection method | Male has control: males mated to female and female dissected/weighed post-insemination, male masturbated via dummy female, natural spawning<br>Male does not have control: Catheter, Males dissected, electroejaculation, males massaged with abdominal pressure |
| Taxonomic Class | Reptilia, Prosomapoda, Monogononta, Mammalia, Malacostraca,<br>Insecta, Hexanauplia, Gastropoda, Clitellata, Chromadorea, Aves, Arachnida, Amphibia, Actinopterygii |
| Population (i.e. Setting) | Laboratory, Domestic, Captive, Wild |
| Method of age estimation | Direct/ Indirect (i.e. inferred from body condition) |
| Trait | Concentration, DNA damage, Ejaculate size, Corrected quantity, Mitochondria, Morphology, Motility, Number of sperm, Oxidative stress, Length, Velocity, Viability |
| Longitudinal sampling | Yes/Semi (when only a subset of males were measured repeatedly)/No |
| Experimental | Yes: manipulated something in addition to male age, or assigned males to specific age groups at the start of the experiment to a target age class<br>No: observational sampling of the available age class distributions opportunistically |
| Gonadosomatic index | 0 to 100 % |
| “Unnatural” manipulations | Yes/No |
| Cold storage of ejaculates (i.e. stored at <5°C) | Yes /No (only for sperm motility, viability, and velocity) |

**Appendix 9:** Formulas used for calculating correlation coefficient (r) from two age groups, multiple age groups, and test statistics/model outputs

A. Two-age groups

To calculate correlation coefficients from studies which reported comparisons between two age groups, we first calculated a standardized mean difference (SMD). SMD here, was calculated using the package *metafor* in R (Viechtbauer, 2010) with the function *escalc.* SMD provides the true strength of an effect by dividing the difference in means between two groups, by their pooled standard deviation. We used the following formula:

$$\text{SMD} = \frac{(Mean\ (old) - Mean\ (young))}{S\ pooled}$$

$$S_{pooled} = \sqrt{\frac{(n_{old} - 1)S_{old}^2 + (n_{young} - 1)S_{young}^2}{n_{old} + n_{young} - 2}}$$

where Mean (o) and (y) and means of the old and young age groups respectively, and S pooled is the pooled standard deviation. These SMD values were then converted to a correlation coefficient using the function *convert_d2r* in the package *meta* (Schwarzer, 2007).

B. Multiple (>2) age groups

To calculate effect sizes from studies which reported means and standard deviations from more than two age groups, we used a simulation (with 1000 iterations). This simulation resulted in a correlation coefficient between the age of males and their means at each age, while weighting the means by their standard deviations. To test for consistency between effect size outcomes from the simulation and the outcomes from SMD, we additionally calculated correlation coefficient for outcomes with only two-groups using the simulation. There was a very strong agreement between r values obtained from these two methods (i.e. SMD and simulation) (R sq.> 0.95).

C. Test statistics (formulae from Koricheva et al, 2013; Polanin and Stiltsveit, 2016)

1. For converting "t" from independent t-test into correlation coefficient (r)

$$r = \frac{t}{\sqrt{(t^2 + degrees\ of\ freedom)}}$$

2. For converting F from ANOVA or ANCOVA with 1 degree of freedom

$$r = \sqrt{(\frac{F}{F + N - 2})}$$

Where N is the sample size of unique number of males

3. For converting spearman's rho to r

$$r = 2 * \sin((\pi * \rho)/6)$$

4. From Mann Whitney-U to r

$$r = \frac{1 - (2 * U)}{n1 * n2}$$

Where n1 and n2 are sample sizes of the younger and older age groups respectively

5. From z-score to r

$$r = \frac{z}{\sqrt{N}}$$

Where N is the sample size of unique number of males

6. For Converting T from Seigel's T test, and converting P values from Mann Whitney U test, to r, the Campbell Collaboration website was used (https://www.campbellcollaboration.org/research-resources/effect-size-calculator.html)
7. For converting R squared and adjusted R squared values to r

$$r = \sqrt{R}^2$$

8. For converting Chisq. values from Chisq. test with one degree of freedom, to r

$$r = \sqrt{(\frac{Chi\ sq.}{N})}$$

**Appendix 10:** Multipliers and signs

If an increase in a trait signified a deleterious effect with increasing age, for example, an increase in sperm abnormal morphology, or sperm DNA damage, we assigned it a multiplier of "-1", whereas if increase in a trait suggested improvement with age, we assigned it a positive multiplier, i.e. "+1". Similarly, when a test statistic was reported (e.g. R sq., correlation coefficient. F values from ANOVA), where older males had worse sperm or ejaculates than younger males, we assigned it a negative multiplier of "-1". Conversely, if older males had better sperm or ejaculates than younger males, we assigned it a multiplier of "+1". Thus, for all the effect sizes in our models, a negative sign indicated reproductive senescence with increasing age, while a positive sign indicated reproductive improvement with increasing age.

**Appendix 11:** Comparison of the three different methods used to calculate effect size: test statistic, standardised mean difference, simulation.

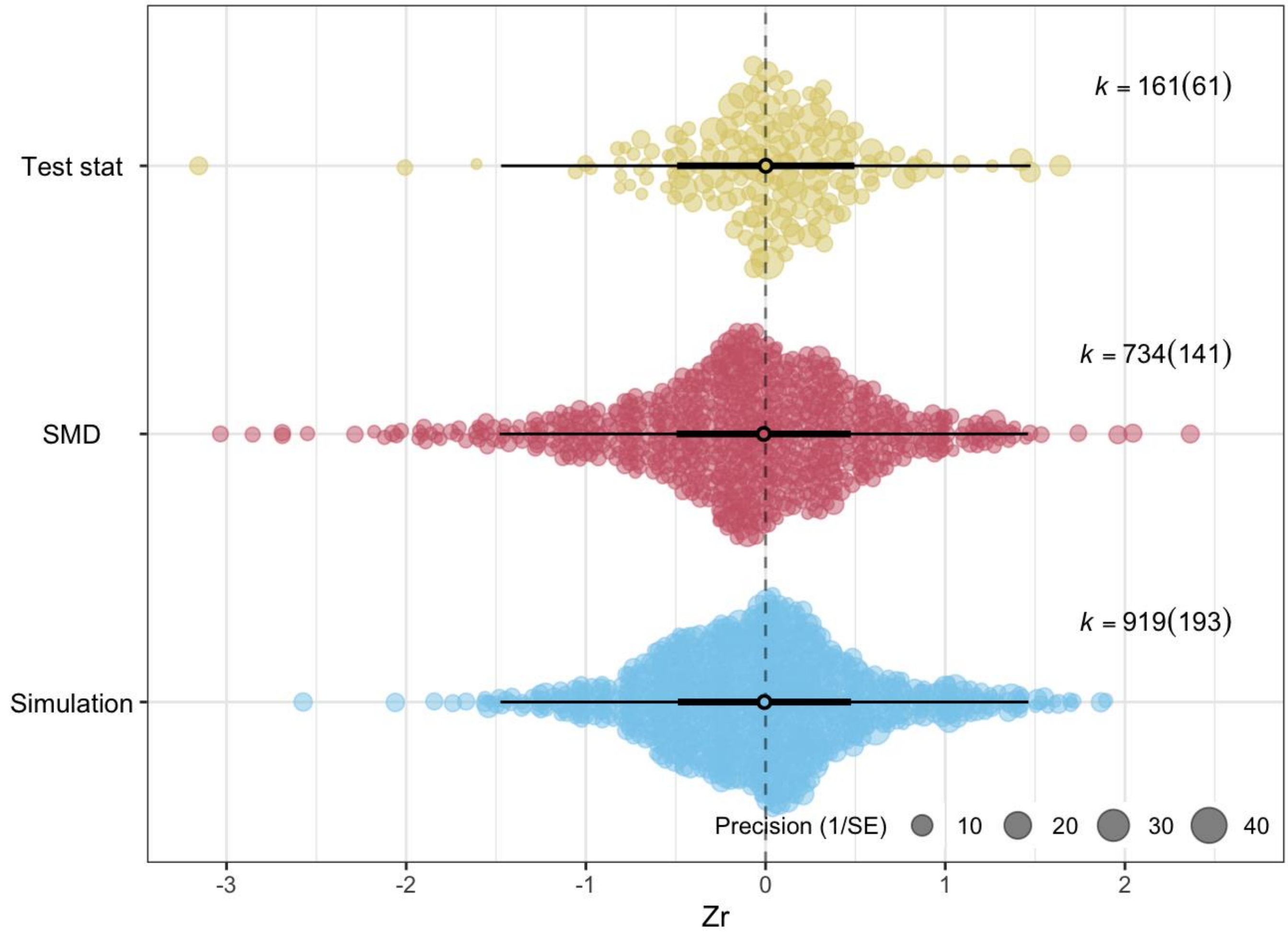


Figure S2. The figure shows no significant difference between the meta-analytical means of the three calculation methods (when calculation method is used as a moderator). The size of each data point represents the precision of the effect size (1/SE). X axis represents values of effect sizes as Fisher's z-transformed correlation coefficient (Zr), while Y axis shows the density distribution of effect sizes. The position of the overall effect is shown by the dark circle, with negative overall values depicting senescence in ejaculates with advancing age while overall positive values showing improvement in ejaculates with advancing male age. Dark error bars (C.I.) show whether overall effect size is significantly different from zero (i.e. not overlapping zero), while light error bars show the prediction interval (P.I.) of effect sizes.
