## Supplementary figures for "No general effects of advancing male age on ejaculates: a meta-analysis across the animal kingdom"

Krish Sangvi and Regina Vega-Trejo

```
knitr::opts_chunk$set(echo = TRUE, fig.align="center", fig.pos = "H", out.extra = "")
```

---

#### Loading packages

---

```
pacman::p_load(tidyverse, metafor, meta, dplyr, kableExtra, GGally,
               bookdown, remotes, ggplot2, Matrix, patchwork, ggpubr,
               rotl, ape, Rcpp, treebase, MCMCglmm, metaviz, matrixcalc,
               orchaRd, #devtools::install_github("daniel1noble/orchaRd", force = TRUE)
               esc, emmeans,
               DHARMa, lme4, lmerTest, ggpointdensity)
```

---

#### Supplementary Figures

---

Data needed for supplementary figures and data manipulation needed to plot figures

```
#import data
spermFinal <- read.csv("spermFinalAllData.csv")
spermFinalFitness<- read.csv("spermFinalFitness.csv")
quadraticData <- read.csv("QuadraticAlldataFinal.csv")

# import phylogenetic matrix
load(file = "./phylo_cor_sperm.Rdata")
load(file = "./phylotree.Rdata")
load(file = "./phyloMCMC.Rdata")
load(file="./phylo_cor_spermFitness.Rdata")
```

```
#Make a new variable for %Lifespan sampled where there are different categories
#Animal_Class = major classes (Insects, Fish, Birds, Mammals) and the rest are classified as 'Others'
#Spell out names for population type

spermFinal<-spermFinal %>%
  mutate(Ls = case_when(LsSampled >=0 & LsSampled <= 25 ~ '0-25%'
                        ,LsSampled >25 & LsSampled <= 50 ~ '25-50%'
                        ,LsSampled >50 & LsSampled <= 75 ~ '50-75%'
                        ,LsSampled >75 & LsSampled <= 100 ~ '75-100%'),
         Animal_Class = case_when(Class == "Actinopterygii" ~ "Actinopterygii"
                                  ,Class == "Aves" ~ "Aves"
                                  ,Class == "Mammalia" ~ "Mammalia"
                                  ,Class == "Insecta" ~ "Insecta"
                                  ,TRUE ~ "Other"),
         Animal_Class = factor(Animal_Class,
                               levels=c("Other", "Mammalia", "Aves", "Actinopterygii", "Insecta")),
         Population = case_when(Population == "C" ~ "Captive",
                                Population == "D" ~ "Domestic",
                                Population == "L" ~ "Lab",
                                Population == "W" ~ "Wild",
                                TRUE ~ Population),
         Population = factor(Population,
                             levels=c("Captive", "Wild", "Lab", "Domestic")),
         Trait = factor(Trait,
                        levels=c("Telomere", "DNA damage", "Oxidative stress",
                                 "Mitochondria", "Sperm length",
                                 "Morphology", "Viability", "Velocity", "Motility",
                                 "Corrected quantity", "Ejaculate size",
                                 "Concentration", "Number of sperm")))
```

#### Prisma diagram

```
knitr::include_graphics("./PrismaAgeSperm.png")
```

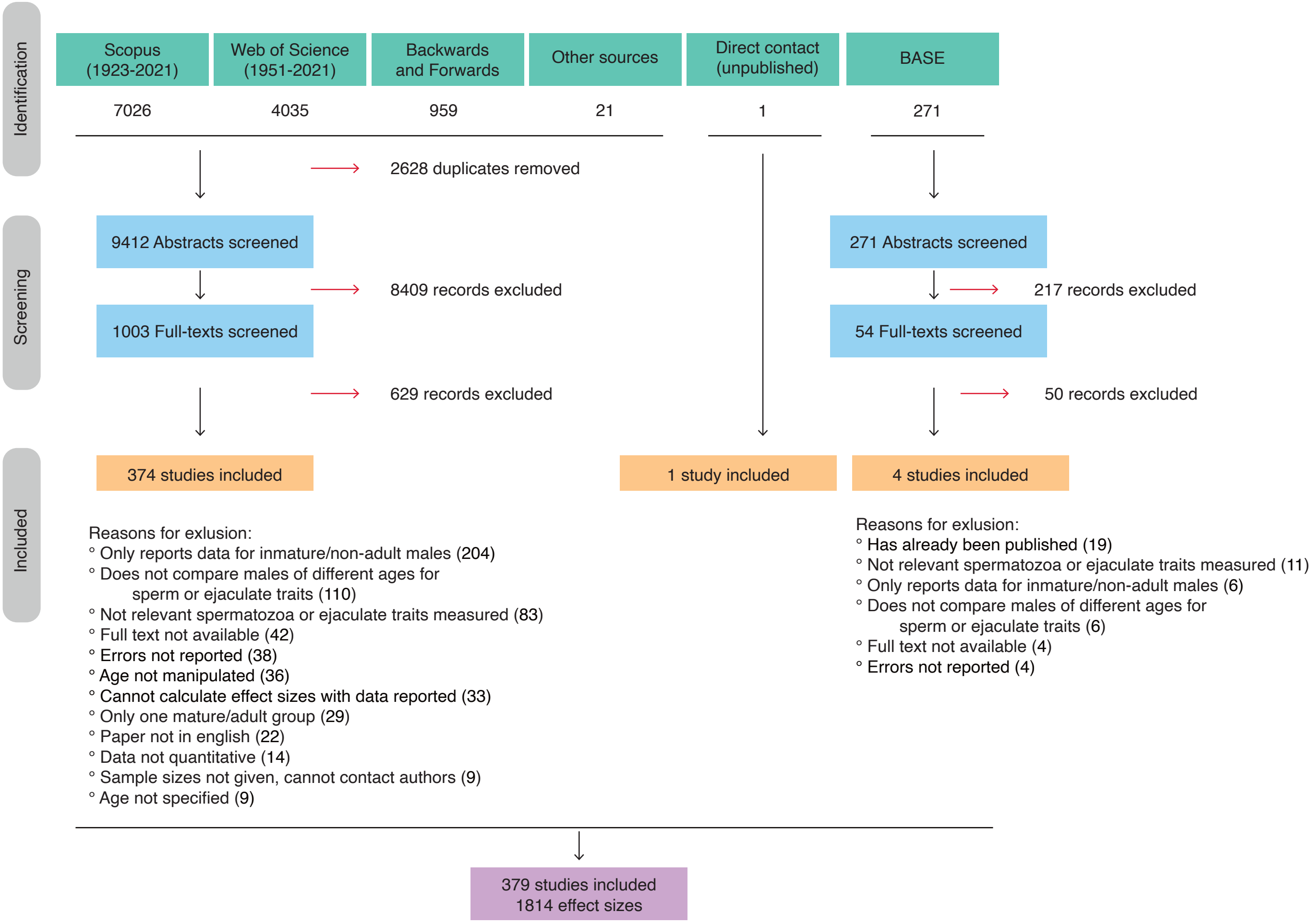


Figure 1: PRISMA diagram describing the search results in different search engines and the different steps of selecting articles for inclusion in the meta-analysis

#### Phylogeny

```r
knitr::include_graphics("./PhyloPlotEdited.png")
```

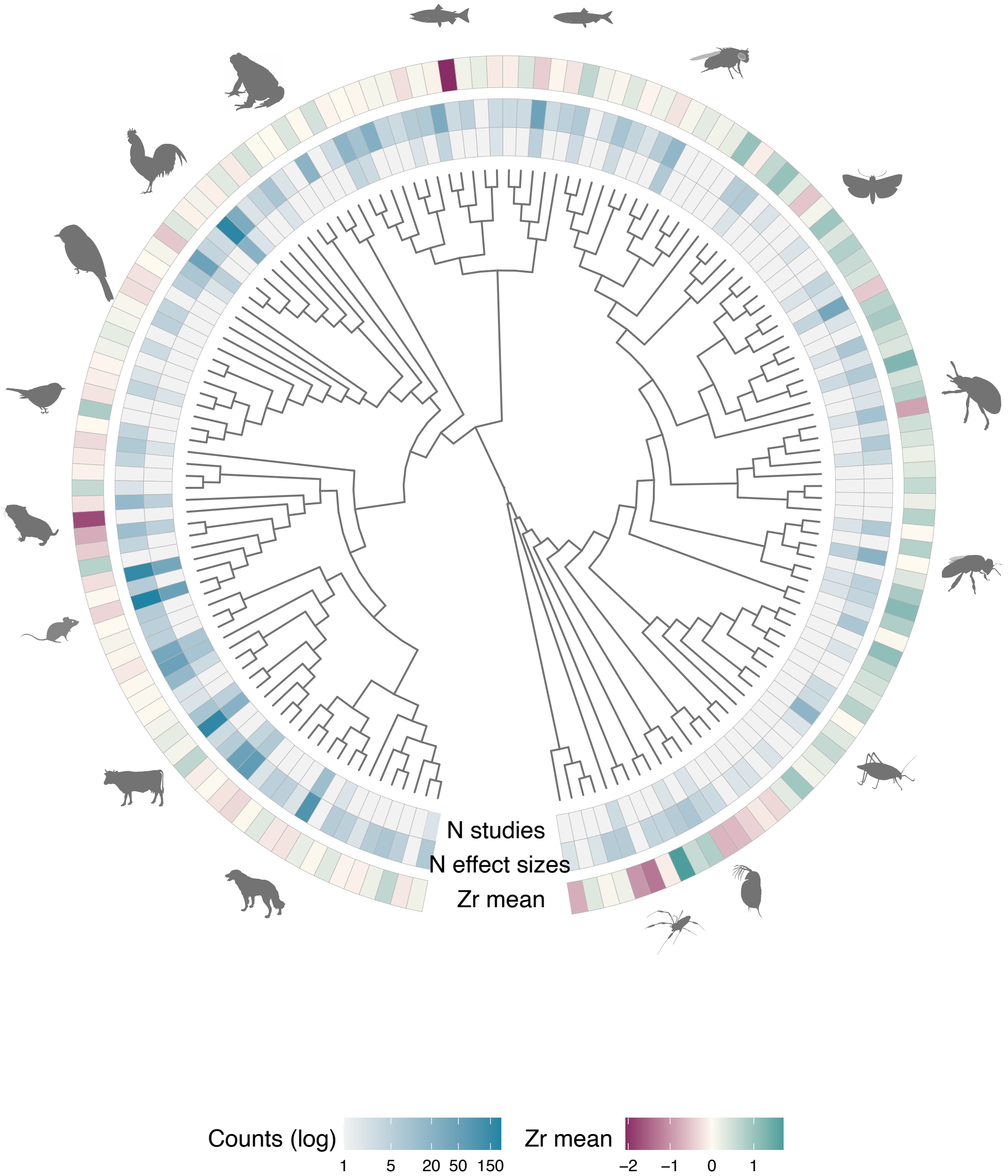


Figure 2: Phylogenetic relatedness explained significant variance in our data ($I^2 = 35.40\%$). Phylogenetic tree of all species (157) included in our meta-analysis, along with the number (N) of studies and effect sizes represented by each species. Overall mean effect size for each species showed as Fisher's z-transformed correlation coefficient (Zr) with negative values representing senescence with increasing age, and positive values representing improvement in ejaculates with increasing age

#### Animal Class

```
load(file = "./Models/Class.model.Rdata")

p.Class<- orchard_plot(Class.model, mod="Class",
                       group = "StudyID",
                       xlab = "Zr",
                       transfm = "none",
                       angle = 0,
                       data = spermFinal)
```

```
p.Class
```

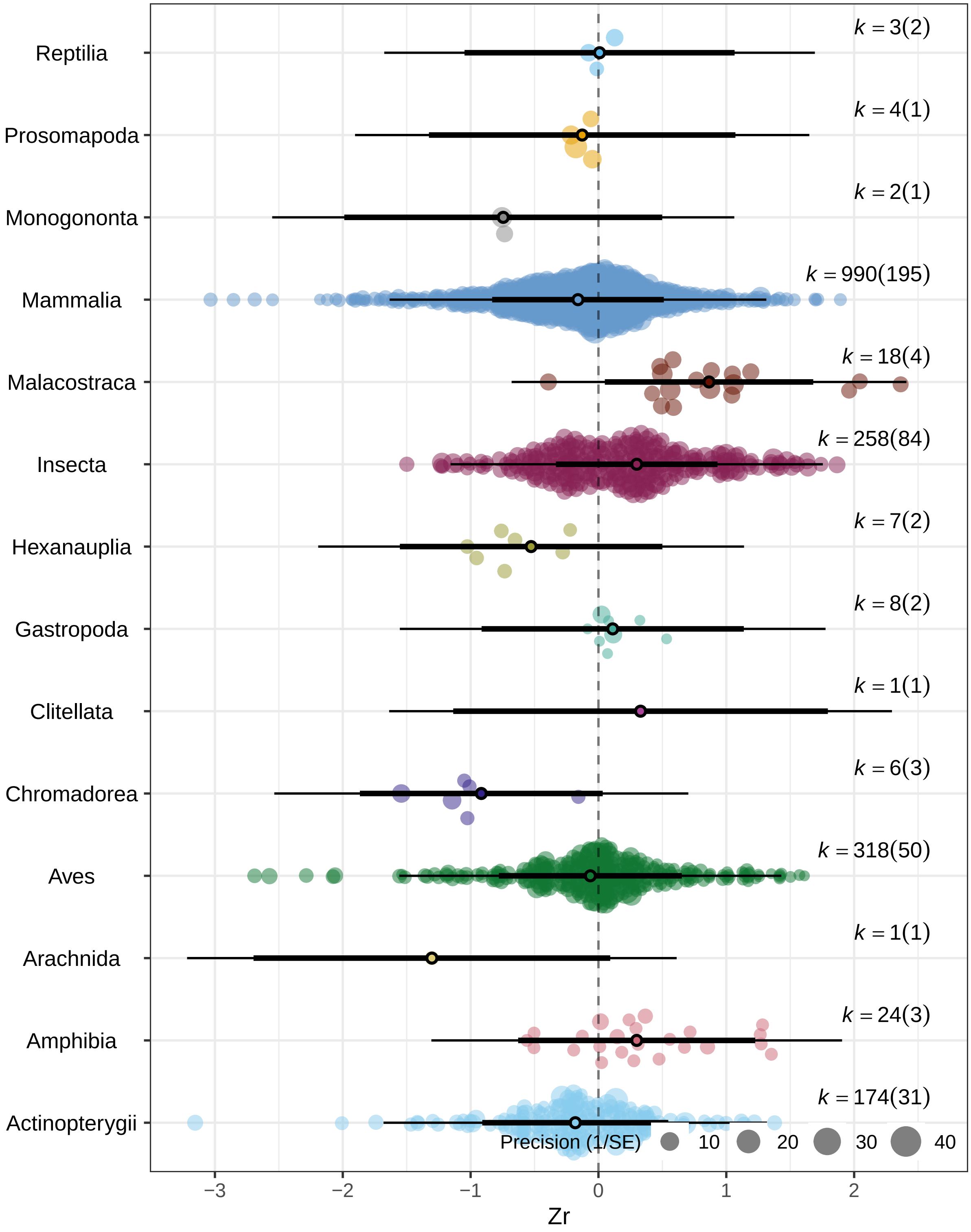


Figure 3: The effect of male age on ejaculates for each class

#### Trait - Major classes

```
#Insects - in main figs
load(file = "./Models/Trait.Insects.model.Rdata")

plot.trait.insects <- orchard_plot(Trait.Insects.model,
                                   group = "StudyID",
                                   mod="Trait",
                                   xlab = "Zr",
                                   transfm = "none",
                                   angle = 0,
                                   data = spermFinal %>% filter(Class %in% c("Insecta"))) +
  scale_color_manual(values=c("#353E7C", "#007094" ,
                              "#008798", "#009B95", "#00BE7D",
                              "#53CC67", "#96D84B", "#CDE030" ,"#FDE333")) +
  scale_fill_manual(values=c("#353E7C", "#007094" ,
                             "#008798", "#009B95", "#00BE7D",
                             "#53CC67", "#96D84B", "#CDE030" ,"#FDE333"))
```

```
## Scale for colour is already present.
## Adding another scale for colour, which will replace the existing scale.
## Scale for fill is already present.
## Adding another scale for fill, which will replace the existing scale.
```

```
##Fish
load(file = "./Models/Trait.Fish.model.Rdata")

p.Trait.Fish.model<- orchard_plot(Trait.Fish.model,
                                  mod="Trait",
                                  group = "StudyID",
                                  xlab = "Zr",
                                  transfm = "none",
                                  data = spermFinal %>% filter(Class %in% c("Actinopterygii"))) +
  theme(axis.text.y = element_text( angle=0)) +
  scale_color_manual(values=c("#46226A", "#353E7C", "#00588B", "#007094" ,
                              "#009B95" ,"#00AE8C", "#00BE7D",
                              "#53CC67", "#96D84B", "#CDE030" ,"#FDE333")) +
  scale_fill_manual(values=c("#46226A", "#353E7C", "#00588B", "#007094" ,
                             "#009B95" ,"#00AE8C", "#00BE7D",
                             "#53CC67", "#96D84B", "#CDE030" ,"#FDE333"))
```

```
## Scale for colour is already present.
## Adding another scale for colour, which will replace the existing scale.
## Scale for fill is already present.
## Adding another scale for fill, which will replace the existing scale.
```

```
##Birds
load(file = "./Models/Trait.Birds.model.Rdata")

p.Trait.Birds.model<- orchard_plot(Trait.Birds.model,
                                   mod="Trait",
                                   group = "StudyID",
```

```r
                                   xlab = "Zr",
                                   transfm = "none",
                                   data = spermFinal %>% filter(Class %in% c("Aves"))) +
  theme(axis.text.y = element_text( angle=0)) +
  scale_color_manual(values=c("#46226A", "#353E7C", "#00588B", "#007094" ,
                              "#008798", "#009B95" ,"#00AE8C", "#00BE7D",
                              "#96D84B", "#CDE030" ,"#FDE333")) +
  scale_fill_manual(values=c("#46226A", "#353E7C", "#00588B", "#007094" ,
                             "#008798", "#009B95" ,"#00AE8C", "#00BE7D",
                             "#96D84B", "#CDE030" ,"#FDE333"))
```

```
## Scale for colour is already present.
## Adding another scale for colour, which will replace the existing scale.
## Scale for fill is already present.
## Adding another scale for fill, which will replace the existing scale.
```

```r
#Mammals
load(file = "./Models/Trait.Mammals.model.Rdata")

p.Trait.Mammals.model<- orchard_plot(Trait.Mammals.model,
                                     mod="Trait",
                                     group = "StudyID",
                                     xlab = "Zr",
                                     transfm = "none",
                                     data = spermFinal %>% filter(Class %in% c("Mammalia"))) +
  theme(axis.text.y = element_text( angle=0)) +
  scale_color_manual(values=c("#4B0055" ,"#46226A", "#353E7C", "#00588B", "#007094" ,
                              "#008798", "#009B95" ,"#00AE8C", "#00BE7D", "#53CC67",
                              "#96D84B", "#CDE030" ,"#FDE333")) +
  scale_fill_manual(values=c("#4B0055" ,"#46226A", "#353E7C", "#00588B", "#007094" ,
                              "#008798", "#009B95" ,"#00AE8C", "#00BE7D", "#53CC67",
                              "#96D84B", "#CDE030" ,"#FDE333"))
```

```
## Scale for colour is already present.
## Adding another scale for colour, which will replace the existing scale.
## Scale for fill is already present.
## Adding another scale for fill, which will replace the existing scale.
```

```r
p.Trait.Fish.model + p.Trait.Birds.model + p.Trait.Mammals.model +
  plot_annotation(tag_levels = "A") &
    theme(plot.tag = element_text(face = 'bold'))
```

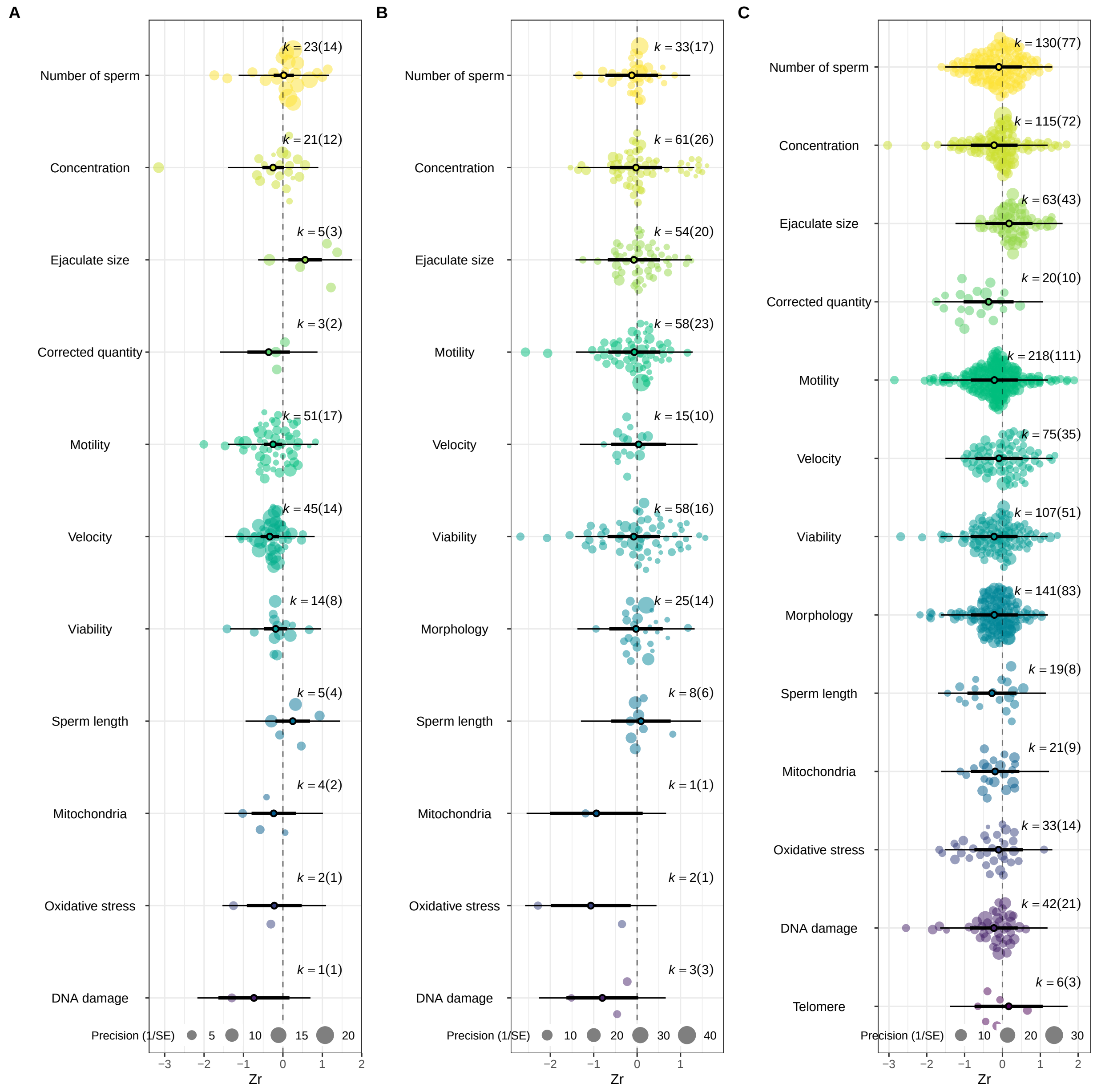


Figure 4: Effect of advancing male age on various ejaculate traits for A. Fish, B. Birds, C. Mammals

#### Trait - Common species

```
load(file = "./Models/Trait.rodents.control.model.Rdata")

p.Trait.rodents.control.model<- orchard_plot(Trait.rodents.control.model,
                                    mod="Trait",
                                    group = "StudyID",
                                    xlab = "Zr",
                                    transfm = "none",
                                    angle = 0,
```

```
                                   data = spermFinal %>%
                                  filter(Manipulation == "No" &
                                           Species %in% c("Mus musculus", "Rattus norvegicus"))) +
  scale_color_manual(values=c("#4B0055" ,"#46226A", "#353E7C", "#007094" ,"#008798",
                              "#009B95" ,"#00AE8C", "#00BE7D", "#53CC67", "#96D84B",
                              "#CDE030" ,"#FDE333"))+
  scale_fill_manual(values=c("#4B0055" ,"#46226A", "#353E7C", "#007094" ,"#008798",
                             "#009B95" ,"#00AE8C", "#00BE7D", "#53CC67", "#96D84B",
                             "#CDE030" ,"#FDE333"))
## Trait - Bulls

load(file = "./Models/Trait.bull.model.Rdata")

p.Trait.bull.model<- orchard_plot(Trait.bull.model,
                                group = "StudyID",
                                mod="Trait",
                                xlab = "Zr",
                                transfm = "none",
                                angle = 0,
                                data = spermFinal %>%
                             filter(Species %in% c("Bos taurus"))) +
  scale_color_manual(values=c("#46226A", "#353E7C", "#00588B",
                              "#008798", "#009B95" ,"#00AE8C", "#00BE7D",
                              "#96D84B", "#CDE030" ,"#FDE333"))+
  scale_fill_manual(values=c("#46226A", "#353E7C", "#00588B",
                             "#008798", "#009B95" ,"#00AE8C", "#00BE7D",
                             "#96D84B", "#CDE030" ,"#FDE333"))

## Trait - chicken
load(file = "./Models/Trait.chicken.model.Rdata")

p.Trait.chicken.model<- orchard_plot(Trait.chicken.model,
                                   mod="Trait",
                                   group = "StudyID",
                                   xlab = "Zr",
                                   transfm = "none",
                                   angle = 0,
                                   data = spermFinal %>%
                                filter(Species %in% c("Gallus gallus"))) +
  scale_color_manual(values=c("#46226A", "#353E7C", "#00588B",
                              "#008798", "#009B95" ,"#00AE8C", "#00BE7D",
                              "#96D84B", "#CDE030" ,"#FDE333"))+
  scale_fill_manual(values=c("#46226A", "#353E7C", "#00588B",
                             "#008798", "#009B95" ,"#00AE8C", "#00BE7D",
                             "#96D84B", "#CDE030" ,"#FDE333"))
```

```
p.Trait.rodents.control.model + p.Trait.bull.model + p.Trait.chicken.model +
  plot_annotation(tag_levels = "A") &
    theme(plot.tag = element_text(face = 'bold'))
```

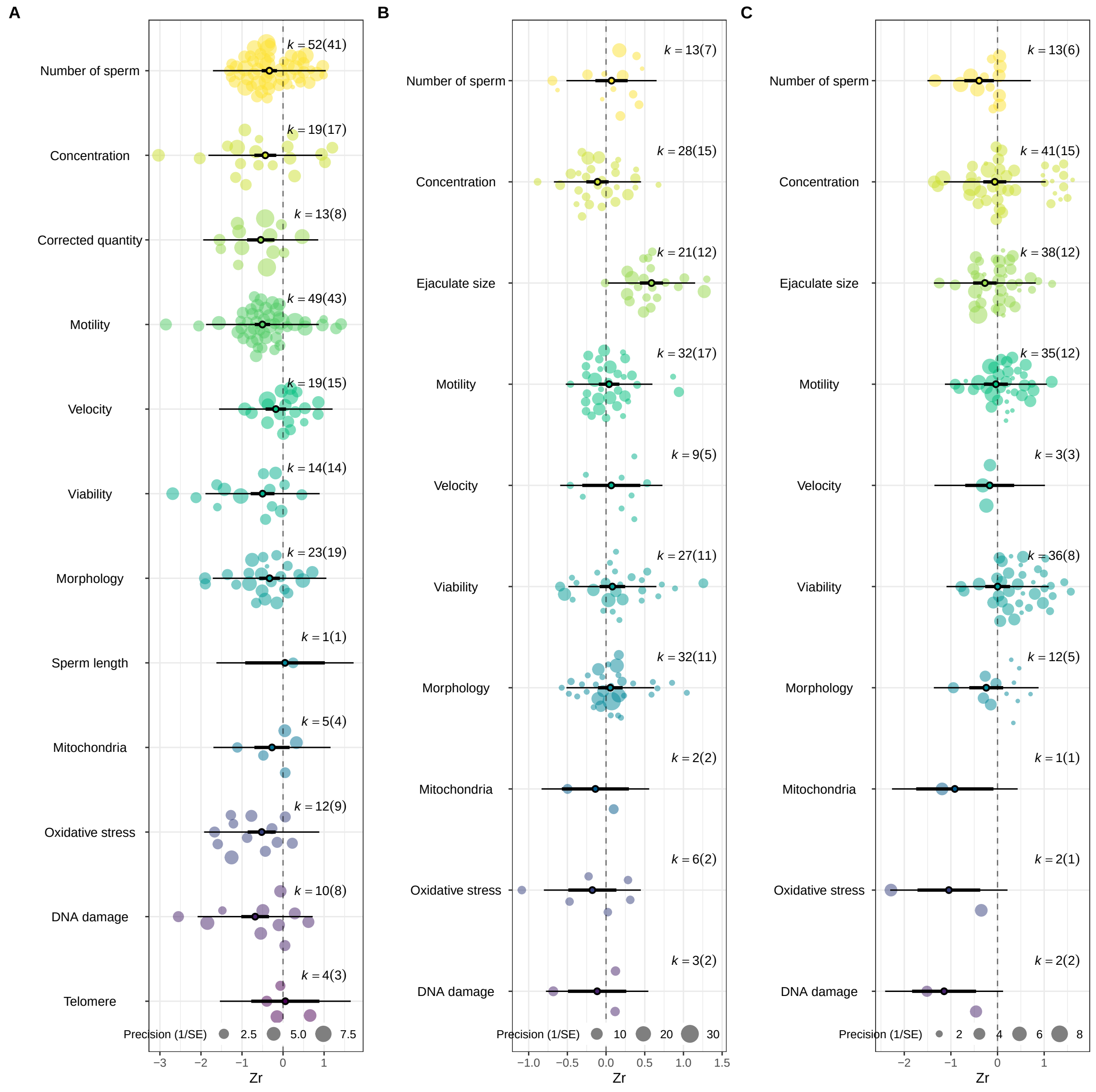


Figure 5: Effect of advancing male age on various ejaculate traits for A. Rodents (*Mus musculus* and *Rattus norvegicus*) that did not go through any manipulation. B. Bulls (*Bos taurus*), C. Chickens (*Gallus gallus*)

### GSI

```
GSIdata <- read.csv("GSIData.csv")
spermFinal <- left_join(spermFinal, GSIdata)
load(file = "./Models/GSI.model.Rdata")

# Plotting by using the residuals of the model
# First making a new data frame which subsets rows which have GSI sampled measurements
#All data
```

```
spermFinal.GSI <- spermFinal[!(is.na(spermFinal$GSI)),]

spermFinal.GSISum <- spermFinal.GSI  %>%
  summarize(N_Studies = length(unique(StudyID)),
            N_EffectSizes = sum(!is.na(esID)))

F.plotting.GSI <- predict(GSI.model)

newdatGSI <- data.frame (GSI = spermFinal.GSI$GSI,
                         Zr = spermFinal.GSI$ZrFinal[!is.na(spermFinal.GSI$ZrFinal)],
                         ZrVar = spermFinal.GSI$VZr[!is.na(spermFinal.GSI$VZr)],
                         fit = F.plotting.GSI$pred, upper = F.plotting.GSI$ci.ub, lower = F.plotting.GSI

newdatGSI$precision <- 1/(sqrt(newdatGSI$ZrVar))
newdatGSI <- newdatGSI[order(newdatGSI$GSI),]

plot.GSI <- ggplot(newdatGSI, aes(x = GSI, y = Zr)) +
  theme(axis.text.y = element_text(size = 12, colour = "black"),
        axis.text.x = element_text(size = 12, colour = "black"),
        panel.background = element_rect(fill = "white"),
        axis.title.y  = element_text(size=15, vjust = 1),
        axis.title.x  = element_text(size=15, vjust = 1),
        panel.border = element_rect(colour = "black", fill=NA, size = 0.1)) +
  #scale_x_continuous(limits = c(-0.5, 27)) +
  #scale_y_continuous(limits = c(-3.2,2.5))+
  geom_point(shape = 21, color = "#03A89E", fill = "#03A89E", alpha = 0.6, cex = 0.2*(newdatGSI$precisio
  geom_hline(yintercept = 0, linetype="dashed", color = "black", size = 0.7) +
  geom_ribbon(aes(ymin = lower, ymax = upper, x = GSI), fill = "black", alpha=0.2) +
  geom_smooth(aes(y = fit, x = GSI), span = 1, color = "black", size = 1) +
  labs(x = "GSI",
       y = "Zr") +
  geom_point(aes(x=14, y= -3), shape = 21, color = "grey65", fill = "grey73", cex = 0.3*5) +
  geom_point(aes(x=16, y= -3), shape = 21, color = "grey65", fill = "grey73", cex = 0.3*15) +
  annotate("text", x = 14, y = 2, size = 3.5, label = "italic(k) == 1508 (288)", parse = TRUE) +
  annotate("text", x = 10, y = -3, size = 3.5, label = "Precision (1/SE)") +
  annotate("text", x = 13, y = -3, size = 3.5, label = "5") +
  annotate("text", x = 15, y = -3, size = 3.5, label = "15")
```

```
plot.GSI
```

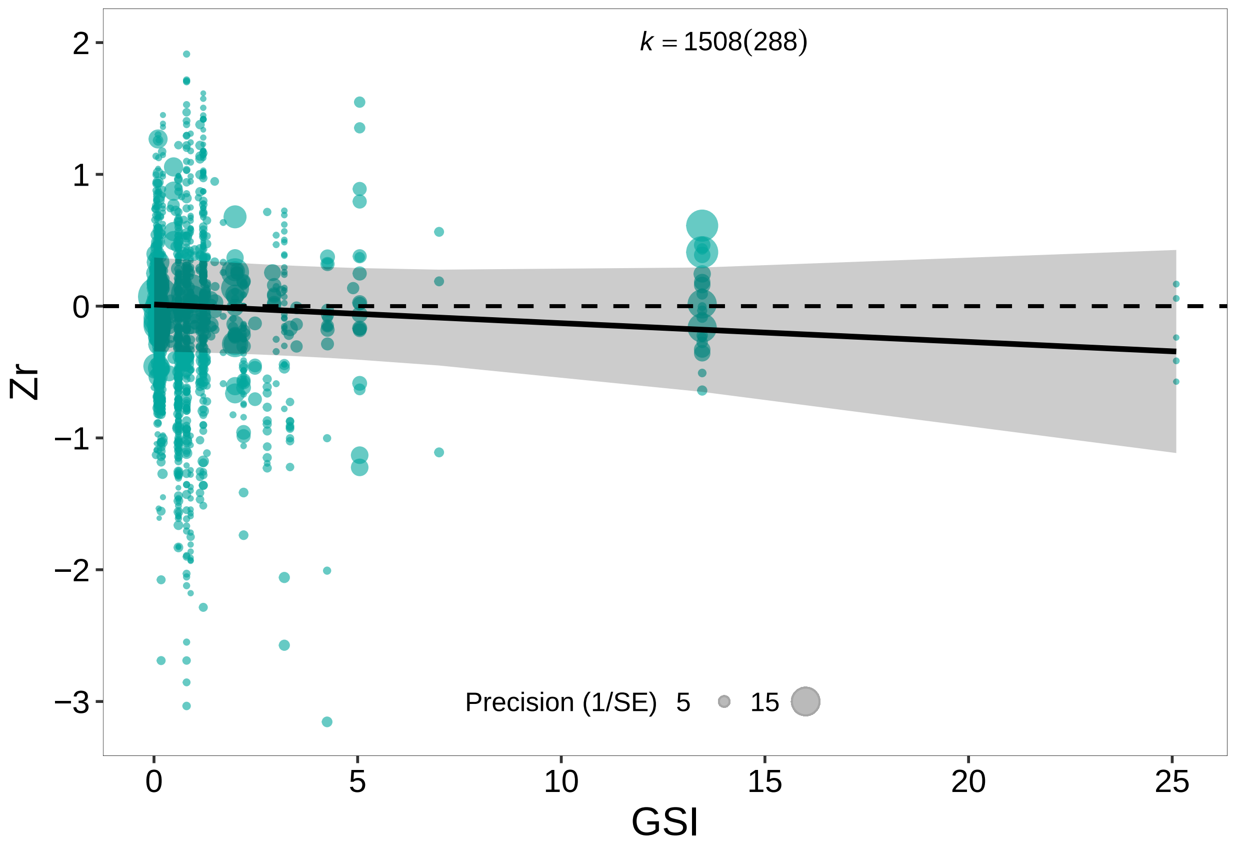


Figure 6: Relationship between gonadosomatic index (GSI) and effect sizes for ejaculate senescence

##### Manipulation

```
load(file = "./Models/ManipulationYN.model.Rdata")

Manipulationdata <- spermFinal %>%
  select(Manipulation, ZrFinal, VZr, StudyID, Cohort, StudyID, esID, Species2, Species) %>%
  drop_na()

p.ManipulalationYN <- orchard_plot(ManipulationYN.model,
             mod="Manipulation",
             group = "StudyID",
             xlab = "Zr",
             transfm = "none",
             angle = 0,
             data = Manipulationdata) +
  scale_color_manual(values=c("#2E5F90","#F7AA2D"))+
  scale_fill_manual(values=c("#2E5F90","#F7AA2D")) +
  scale_x_discrete(labels=c("Yes" = "Manipulated", "No" = "Control"))

#p.ManipulalationYN
```

```r
#Note that this only includes data for manipulated males only
#Let's group type of manipulation into categories that are easier to understand
# Toxin/chemical and medicine into pharmacological intervention

spermFinal <- spermFinal %>%
  mutate(ManipulationType2 = case_when(ManipulationType == "Toxin/chemical" ~ "Pharmacological interven
                                ManipulationType == "Medicine" ~ "Pharmacological intervention",
                                ManipulationType == "Light" ~ "Light regime",
                                TRUE ~ ManipulationType))

load(file = "./Models/ManipulatedType.model.Rdata")
manipulated.dat <- spermFinal %>% filter(ControlOrTreatment %in% c("Treatment"))

p.ManipulatedType.model<- orchard_plot(ManipulatedType.model,
                                       mod="ManipulationType2",
                                       xlab = "Zr",
                                       transfm = "none",
                                       group = "StudyID",
                                       angle = 0,
                                       data = manipulated.dat) +
  scale_color_manual(values=c("#7F3C8D", "#11A579", "#3969AC", "#F2B701",
                              "#E73F74", "#80BA5A", "#E68310", "#008695")) +
  scale_fill_manual(values=c("#7F3C8D", "#11A579", "#3969AC", "#F2B701",
                             "#E73F74", "#80BA5A", "#E68310", "#008695"))
#p.ManipulatedType.model
```

```r
p.ManipulalationYN / p.ManipulatedType.model +
  plot_annotation(tag_levels = "A") &
    theme(plot.tag = element_text(face = 'bold'))
```

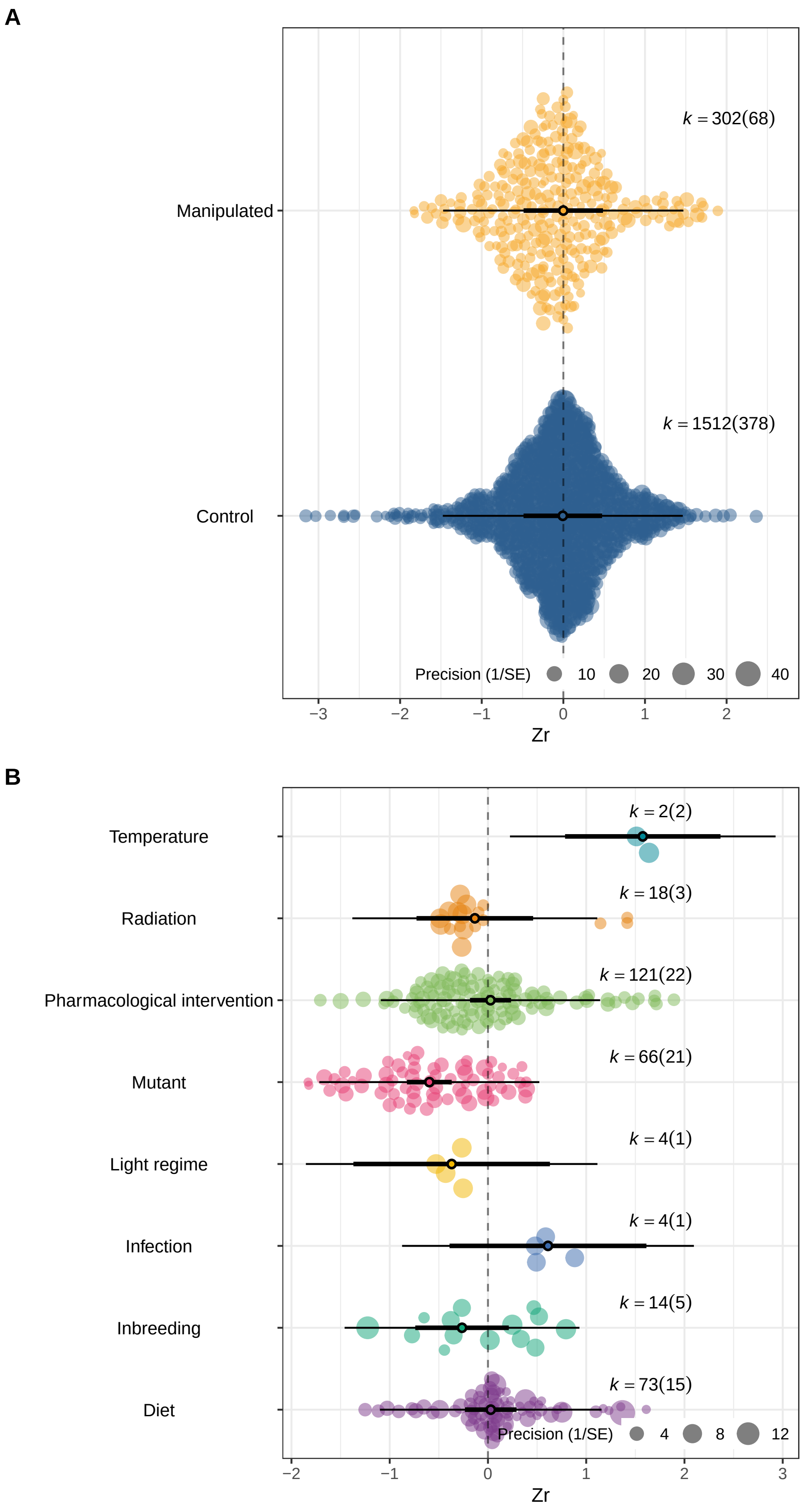


Figure 7: A. Effect of advancing male age on ejaculates for studies with 'control' vs 'manipulated' males, B. Effects of advancing male age on ejaculates for each type of manipulation/treatment including manipulated males only

#### Sampling method

```r
load(file = "./Models/Longi.model.Rdata")

p.Longi.model<- orchard_plot(Longi.model,
                             mod="Longitudinal",
                             group = "StudyID",
                             xlab = "Zr",
                             transfm = "none",
                             angle = 0,
                             data = spermFinal) +
  scale_x_discrete(labels=c("Yes" = "Longitudinal",
  "Semi" = "Semi-longitudinal", "No" = "Cross-sectional"))
```

```r
p.Longi.model
```

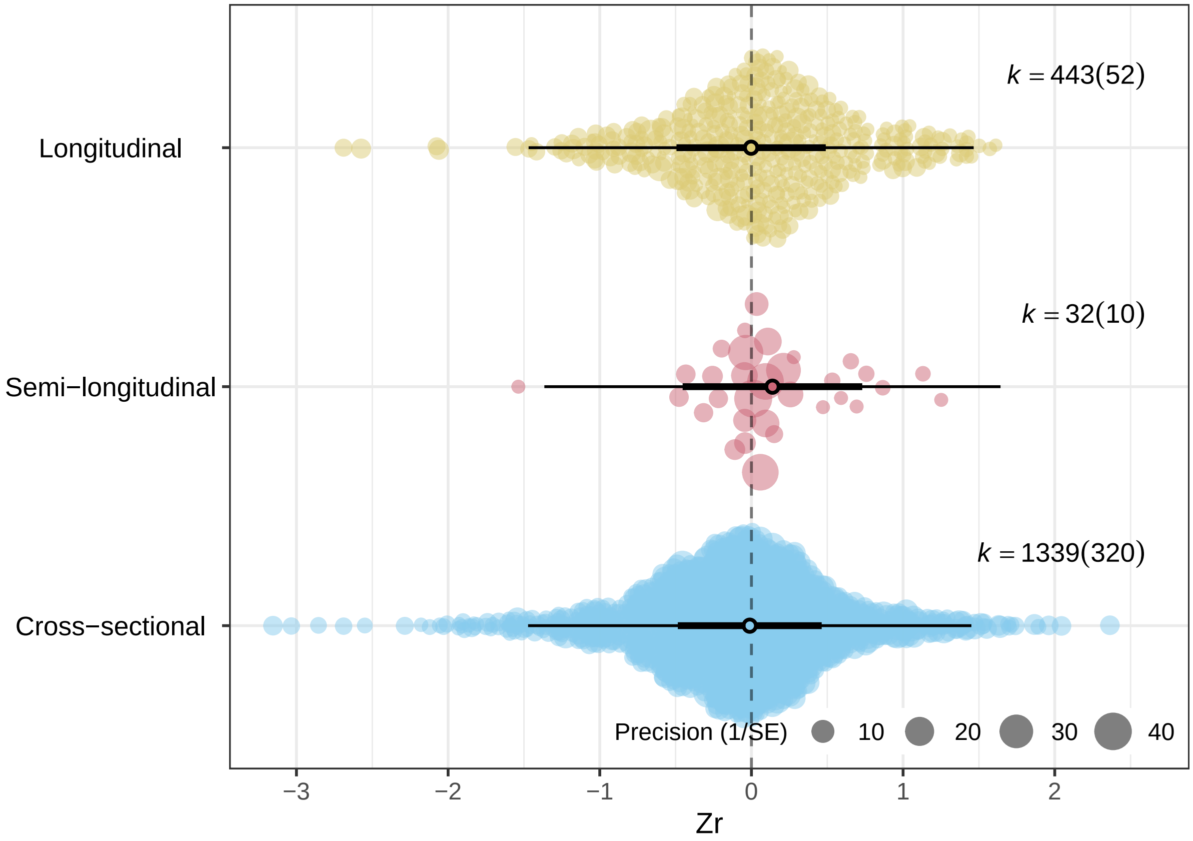


Figure 8: Effect of advancing male age on ejaculates for studies with longitudinal vs cross-sectional sampling on males

#### Ejaculate collection method

```
#yes vs no
load(file = "./Models/ejacControl.model.Rdata")

spermFinal.extract<-spermFinal %>%
  filter(ControlOverEjaculation %in% c("y" , "n"))

p.ejacControl<- orchard_plot(ejacControl.model,
                             mod="ControlOverEjaculation",
                             group = "StudyID",
                             xlab = "Zr",
                             transfm = "none",
                             angle = 0,
                             data = spermFinal.extract) +
  scale_x_discrete(labels = c("Y" = "Control over ejaculation", "N" = "No control over ejaculation"))

load(file = "./Models/Extraction.model.Rdata")

#Note that some effect sizes didn't mention what method was used for what late collection, therefore be
spermFinal.collection <- spermFinal %>%
  filter(ExtractionMethod != "" )

p.Extraction.model<- orchard_plot(Extraction.model,
                                  mod="ExtractionMethod",
                                  group = "StudyID",
                                  xlab = "Zr",
                                  transfm = "none",
                                  angle = 0,
                                  data = spermFinal.collection)
#p.Extraction.model
```

```
p.ejacControl / p.Extraction.model +
  plot_annotation(tag_levels = "A") &
    theme(plot.tag = element_text(face = 'bold'))
```

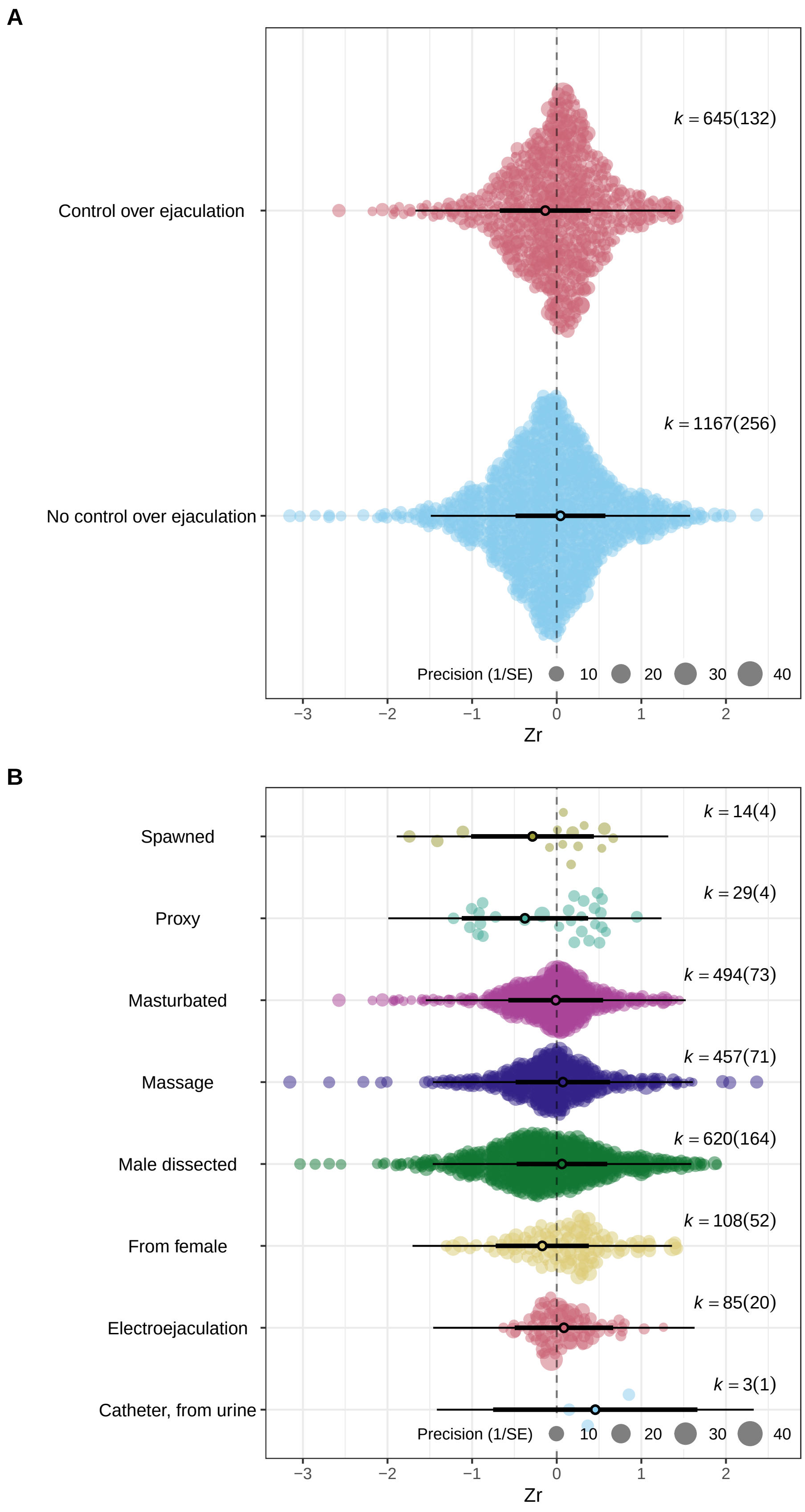


Figure 9: A. Effect of male's 'control' over ejaculation on advancing male age. B. Effect of advancing male age on ejaculates for each type of ejaculate collection method

#### Lifespan Sampled

```
ClassLSsummary <- spermFinal %>% group_by(StudyID, Animal_Class) %>%
  summarise(LsSampledmean =mean(LsSampled))

Class_LS2 <- ggplot(ClassLSsummary,
                   aes (x=fct_rev(Animal_Class), y= LsSampledmean, fill = fct_rev(Animal_Class))) +
  geom_boxplot(outlier.shape = NA) +
  scale_fill_manual(values=c("#26A63A" , "#9BB306" ,"#E1BB4E", "#FFC59E" ,"#BFBFBF")) +
  geom_jitter(size=0.1, alpha = 0.5) +
  theme_classic() +
  xlab("Animal Class") +
  ylab("Mean lifespan sampled") +
  theme(legend.position="none",
        text=element_text(size=16))

PopLSsummary <- spermFinal %>% group_by(StudyID, Population) %>%
  summarise(LsSampledmean =mean(LsSampled))

Pop_LS2 <- ggplot(PopLSsummary, aes (x=fct_rev(Population), y= LsSampledmean, fill = fct_rev(Population
  geom_boxplot(outlier.shape = NA) +
  scale_fill_manual(values=c("#E16462FF", "#FCA636FF", "#B12A90FF", "#6A00A8FF")) +
  geom_jitter(size=0.1, alpha = 0.5) +
  theme_classic() +
  xlab("Population type") +
  ylab("Mean lifespan sampled") +
  theme(legend.position="none",
        text=element_text(size=16))

Spp_LSsummary <- spermFinal %>%
  filter(Species %in% c("Bos taurus", "Rattus norvegicus", "Mus musculus", "Gallus gallus"))%>%  group_
  summarise(LsSampledmean =mean(LsSampled))

Spp_LS2 <- ggplot (data = Spp_LSsummary, aes (x=factor(Species, level = c("Gallus gallus", "Rattus norv
  geom_boxplot(outlier.shape = NA) +
  scale_fill_manual(values=c("#A4473D", "#A57E00", "#86B13E", "#53DCAB" )) +
  geom_jitter(size=0.1) +
  theme_classic() +
  xlab("Species") +
  ylab("Mean lifespan sampled") +
  theme(legend.position="none",
        text=element_text(size=16))
```

```
(Pop_LS2 / Spp_LS2) + Class_LS2 + plot_annotation(tag_levels = "A") &
    theme(plot.tag = element_text(face = 'bold'))
```

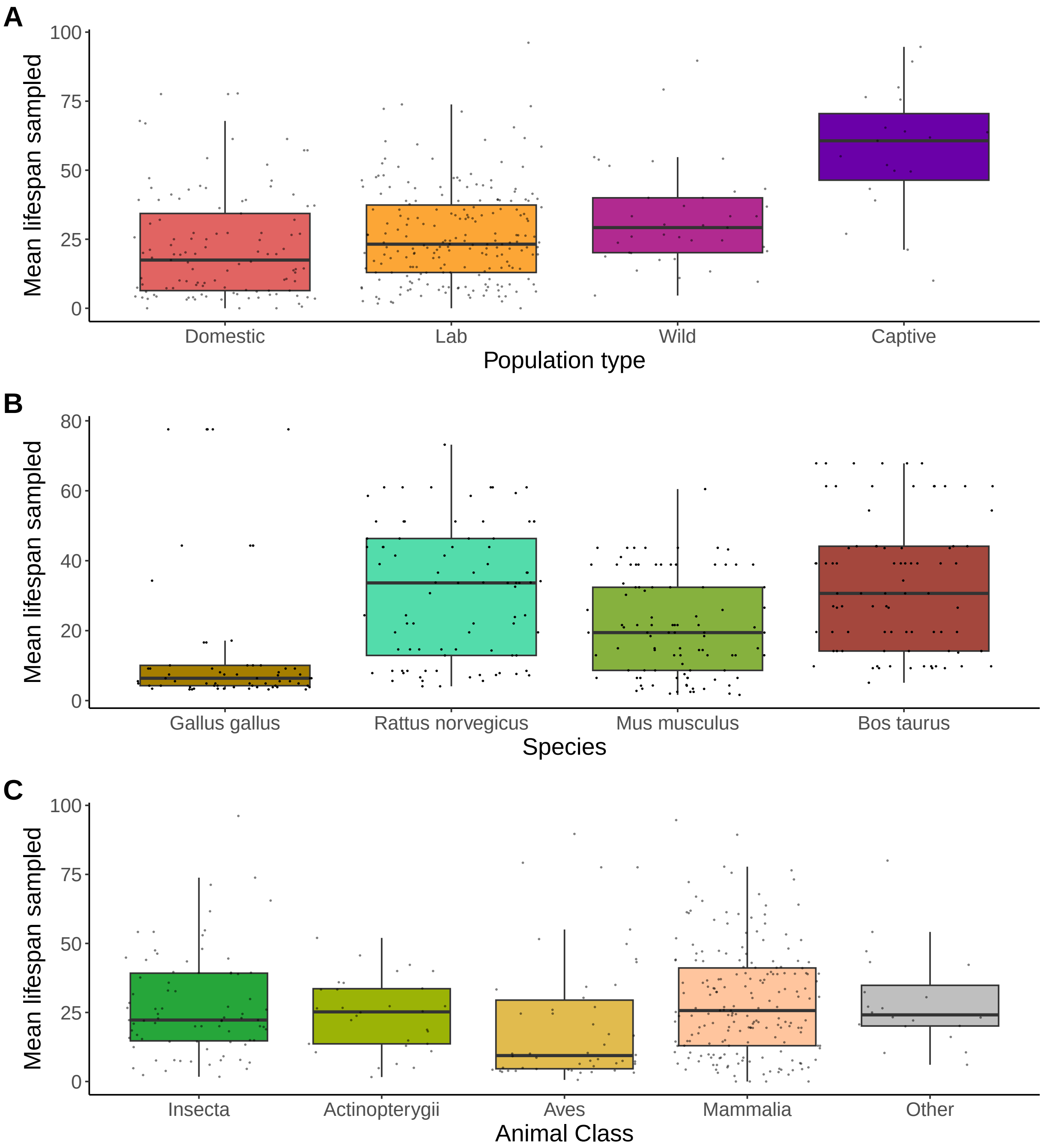


Figure 10: Medians (50%) and interquartile ranges (5%, 25%, 75%, 95%) of proportion of maximum adult lifespan sampled for A. Different population types, B. Four most common species in our dataset, and C. Different Animal Classes (note that animal classes with less than 25 effect sizes were grouped together in 'Other'). Each point refers to the average lifespan sampled from a study

##### Cold storage

```
load(file = "./Models/ColdStorage.model.Rdata")

Storagedata <- spermFinal %>%
  filter(Trait  %in% c("Motility", "Velocity", "Viability") & ColdStorage %in% c("N", "Y"))

p.ColdStorage.model<- orchard_plot(ColdStorage.model,
                             mod="ColdStorage",
                             group = "StudyID",
                             xlab = "Zr",
                             transfm = "none",
                             angle = 0,
                             data = Storagedata) +
  scale_color_manual(values=c("#A4473D", "#86B13E")) +
  scale_fill_manual(values=c("#A4473D", "#86B13E")) +
  scale_x_discrete(labels = c("Y" = "Yes", "N" = "No"))
```

```
p.ColdStorage.model
```

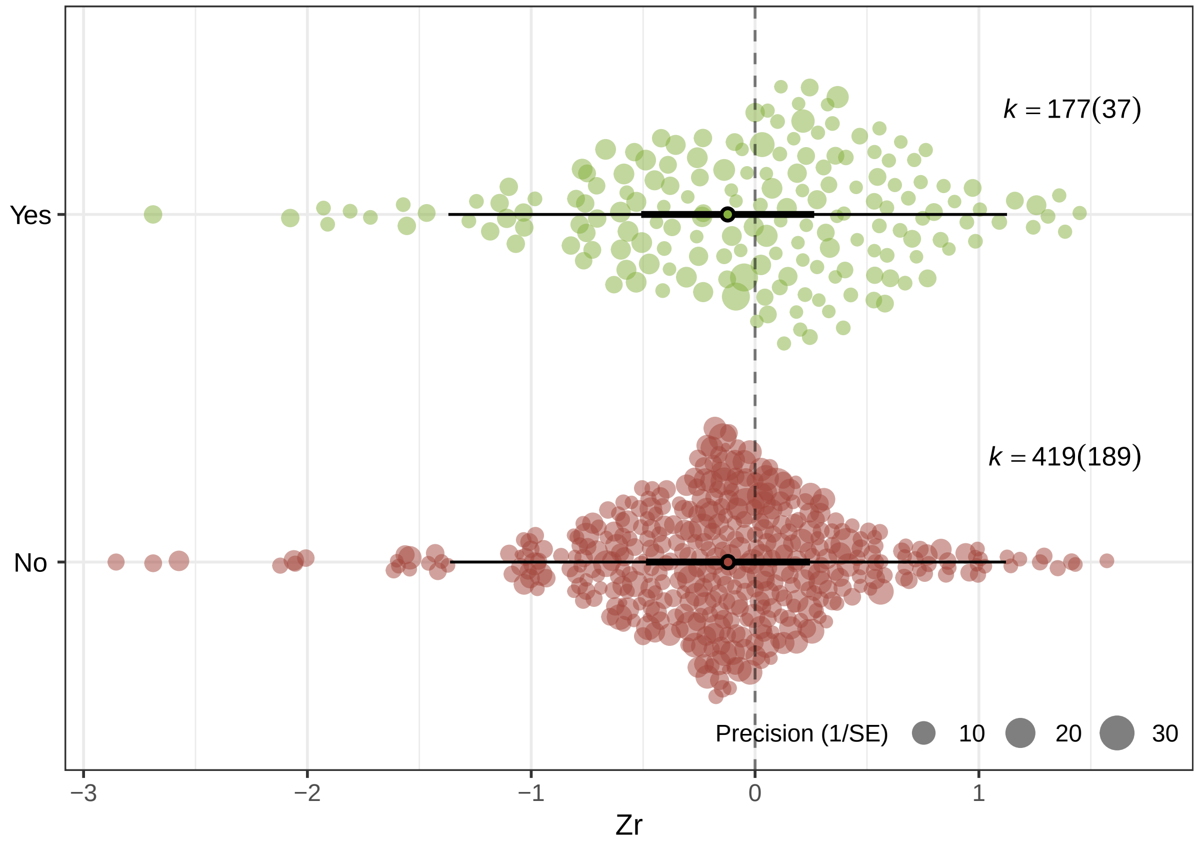


Figure 11: Effect of advancing male age on ejaculates for studies with cold storage (yes vs no) for motility, velocity, and viability

#### Reproductive output

```
load(file = "./Models/Fitness.model.Rdata")

p.Fitness.model <- orchard_plot(Fitness.model,
                                mod = "FitnessOrEjaculate",
                                group = "StudyID",
                                xlab = "Zr",
                                transfm = "none",
                                angle = 0,
                                data = spermFinalFitness) +
                                scale_x_discrete(labels = c("Fitness" = "Reproductive output", "Ejacula

#p.Fitness.model

load(file = "./Models/Fitness.Trait.model.Rdata")

p.Fitness.Trait.model <- orchard_plot(Fitness.Trait.model ,
                                      mod="Trait",
                                      group = "StudyID",
                                      xlab = "Zr",
                                      transfm = "none",
                                      angle = 0,
                                      data = spermFinalFitness %>%
                                        filter(FitnessOrEjaculate %in% c("Fitness"))) +
  scale_color_manual(values=c("#CC0000", "#006600")) +
  scale_fill_manual(values=c("#CC0000", "#006600"))
```

```
## Scale for colour is already present.
## Adding another scale for colour, which will replace the existing scale.
## Scale for fill is already present.
## Adding another scale for fill, which will replace the existing scale.
```

```
#p.Fitness.Trait.model
```

```
p.Fitness.Trait.model / p.Fitness.model  +
  plot_annotation(tag_levels = "A") &
    theme(plot.tag = element_text(face = 'bold'))
```

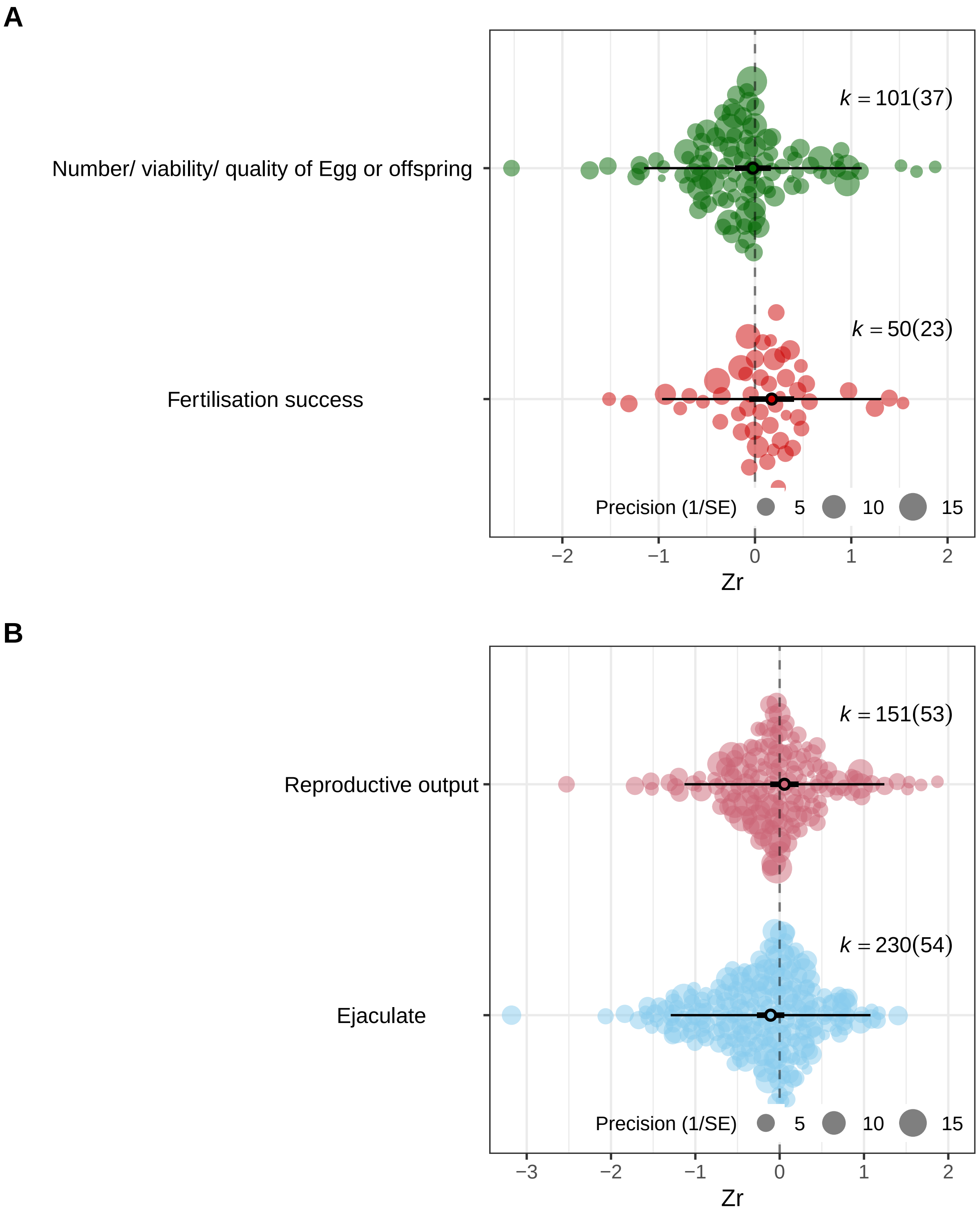


Figure 12: A. Effect of advancing male age on type of fitness trait measured in unmanipulated males. B. Effects of advancing male age on reproductive output and on ejaculates from studies which measure both traits

#### Quadratic plots

```
p.quadratic.trait <- ggplot(quadraticData,
                            aes (x= Standardised_age, y= Standardised_trait, colour=Trait)) +
  geom_point (aes(size=log.N_males), shape = 1, alpha = 0.6) +
  stat_smooth(aes(y = Standardised_trait, fill=Trait), method = "lm",
              formula = y ~ x + I(x^2), size = 1.5) +
  scale_color_manual(values=c("#B65A49", "#914CB1", "#97C160")) +
  scale_fill_manual(values=c("#B65A49", "#914CB1", "#97C160")) +
  xlab("Standardised male age") +
  ylab("Standardised ejaculate trait") +
  theme_classic()
```

```
p.quadratic.trait
```

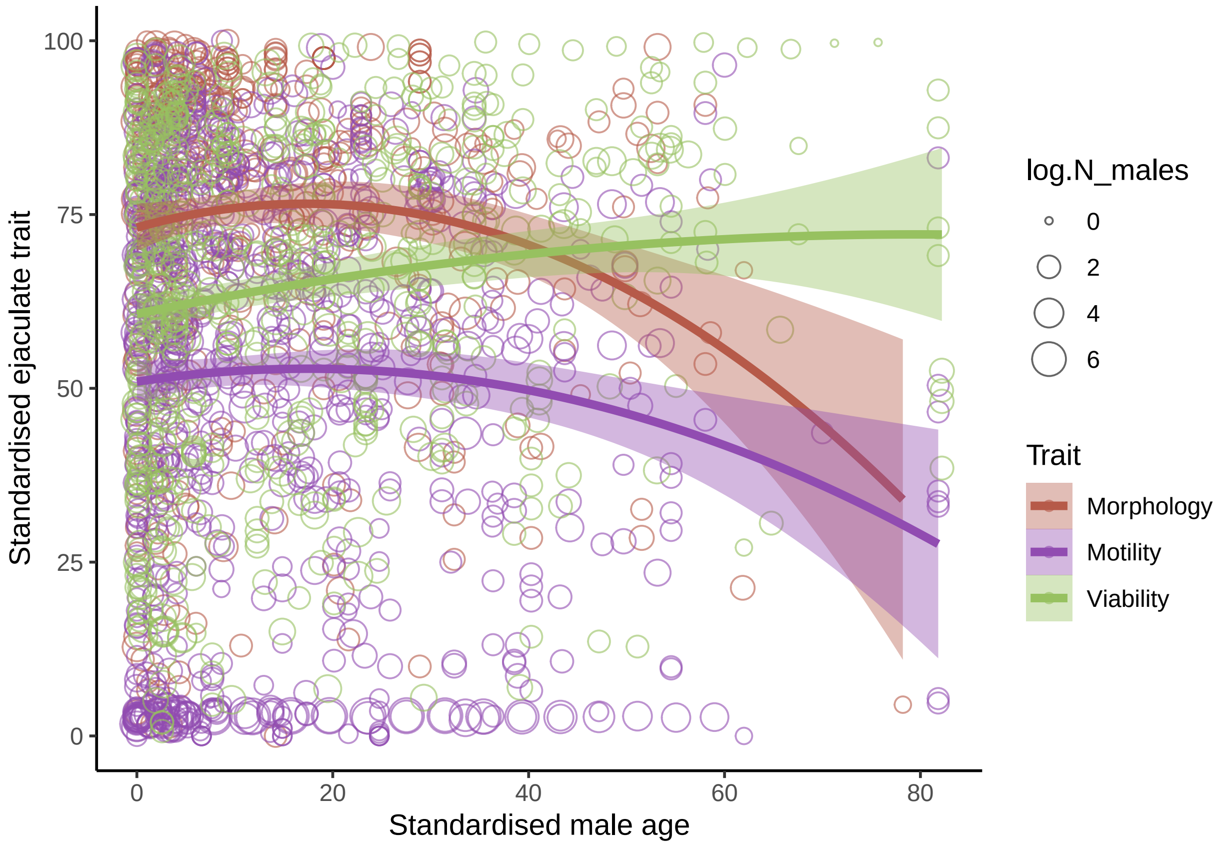


Figure 13: Effects of standardised male age (X axis) on standardised ejaculate trait values (separately shown for the three traits of % morphologically normal, % motile, and % viable sperm). Standardised male age calculated as proportion of maximum adult lifespan represented by the specific age class. Standardised trait value calculated as percentage of with morphologically normal (N= 85 studies, k= 153 effect sizes), viable (N= 81 studies, k= 193 effect sizes), or motile (N= 137 studies, k= 294 effect sizes) sperm

##### Funnel plot

```r
load(file = "./Models/null.model.Alldata.Rdata")

f.null <- funnel(null.model.Alldata, level=c(90, 95, 99),
                 shade=c("white", "gray55", "gray75"),
                 yaxis="seinv", refline=0,
                 pch=1,
                 cex= 0.4, bg= "white",
                 legend=FALSE)
```

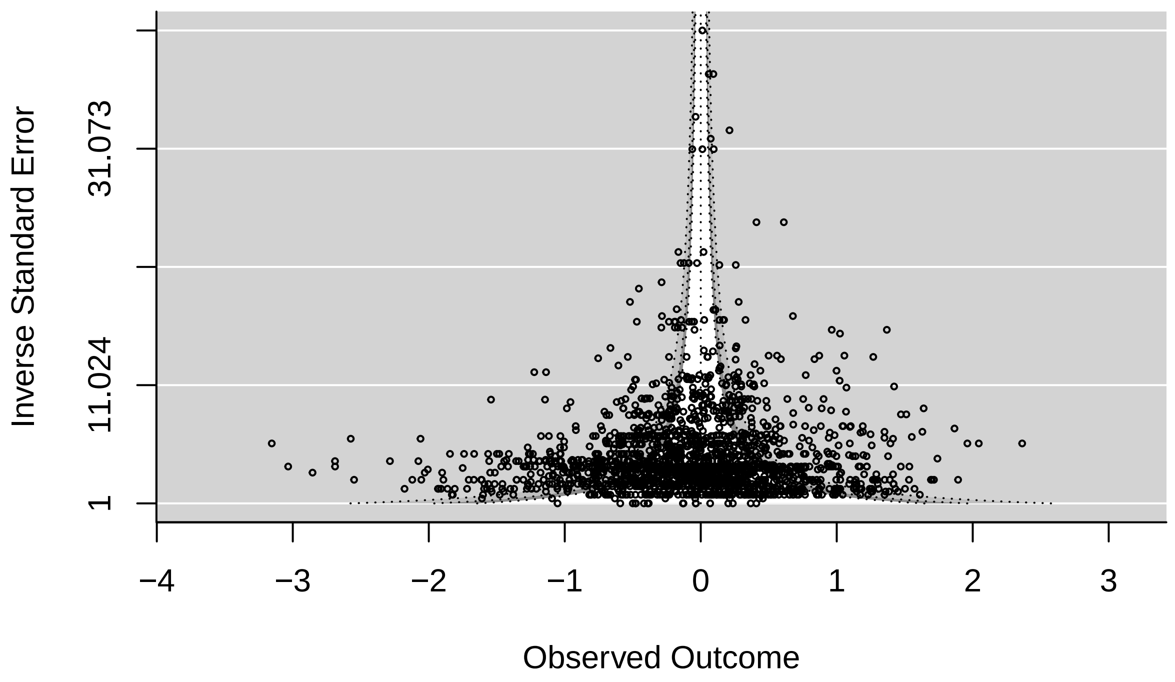


Figure 14: Funnel plot with precision as a measure of uncertainty to explore the existence of outliers in the meta-analytic dataset

##### Small-study and time-lag bias

```r
load(file = "./Models/publication.bias.model.r.all.Rdata")

spermFinalYears <- spermFinal %>%
  filter(Year %in% (1969:2021))

spermFinalYears$Year <- as.numeric(spermFinalYears$Year) # converting year to number from factor
```

```r
spermFinalYears$sei <- sqrt(spermFinalYears$VZr) #calculate sei for this dataset

# mean-centering year of publication to help with interpretation. "0" is the mean of all years
spermFinalYears$year.c <- as.vector(scale(spermFinalYears$Year, scale = F))

## Small-study bias
predict.publication.bias.model.r.all.se.plot.1 <- predict(publication.bias.model.r.all,
    newmods = cbind(seq(min(spermFinalYears$sei), max(spermFinalYears$sei), 0.005), c(0)))

newdatsei <- data.frame(sei = seq(min(spermFinalYears$sei),
                              max(spermFinalYears$sei), 0.005),
                     fit = predict.publication.bias.model.r.all.se.plot.1$pred,
                     upper = predict.publication.bias.model.r.all.se.plot.1$ci.ub,
                     lower = predict.publication.bias.model.r.all.se.plot.1$ci.lb,
                     stringsAsFactors = FALSE)

ZrFinal <- newdatsei$fit

spermFinalYears$precision <- 1/(sqrt(spermFinalYears$VZr))

plotSmallStudybias <- ggplot(data=spermFinalYears, aes(x = sei, y = ZrFinal)) +
  theme(axis.text.y = element_text(colour = "black"),
        axis.text.x = element_text(colour = "black"),
        panel.background = element_rect(fill = "white"),
        axis.title.y  = element_text(vjust = 1),
        axis.title.x  = element_text(vjust = 1),
        panel.border = element_rect(colour = "black", fill=NA, size = 0.5)) +
  geom_point(shape = 21, color = "black", fill = "#458B00", alpha = 0.3, cex = 0.4*(spermFinalYears$pre
  geom_ribbon(data=newdatsei, aes(ymin = lower, ymax = upper, x = sei), fill = "black", alpha=0.1) +
  geom_smooth(data=newdatsei, aes(y = fit, x = sei), span = 1, color = "black", size = 1) +
  geom_hline(yintercept=0, linetype="dashed", color = "black") +
    labs(x = "Standard error (SE)",
       y = "Zr") +
  geom_point(aes(x=0.8, y= -3), shape = 21, color = "grey65", fill = "grey73", cex = 0.4*15) +
  geom_point(aes(x=0.72, y= -3), shape = 21, color = "grey65", fill = "grey73", cex = 0.4*5) +
  annotate("text", x = 0.55, y = -3, label = "Precision (1/SE)") +
  annotate("text", x = 0.75, y = -3, label = "5") +
  annotate("text", x = 0.85, y = -3, label = "15")+
  theme(text =element_text(size = 15))

## Time-lag bias

predict.publication.bias.model.r.all.se.plot.2 <- predict(publication.bias.model.r.all,
    newmods = cbind(mean(spermFinalYears$sei), seq(min(spermFinalYears$year.c), max(spermFinalYears$yea
        0.25)))

newdatYears <- data.frame(year = seq(min(spermFinalYears$Year),
                              max(spermFinalYears$Year), 0.25),
                     fit = predict.publication.bias.model.r.all.se.plot.2$pred,
                     upper = predict.publication.bias.model.r.all.se.plot.2$ci.ub,
                     lower = predict.publication.bias.model.r.all.se.plot.2$ci.lb,
                     stringsAsFactors = FALSE)
```

```
ZrFinal2 <- newdatYears$fit
#This overwrites the above so let's make a new variable
spermFinalYears <- spermFinalYears %>%
  mutate(ZrFinal2 = ZrFinal)
spermFinalYears$precision2 <- 1/(sqrt(spermFinalYears$VZr))

plotYearbias <- ggplot(data=spermFinalYears, aes(x = Year, y = ZrFinal2)) +
  theme(axis.text.y = element_text(colour = "black"),
        axis.text.x = element_text(colour = "black"),
        panel.background = element_rect(fill = "white"),
        axis.title.y  = element_text(vjust = 1),
        axis.title.x  = element_text(vjust = 1),
        panel.border = element_rect(colour = "black", fill=NA, size = 0.5)) +
  geom_point(shape = 21, color = "black", fill = "#4876FF", alpha = 0.3, cex = 0.4*(spermFinalYears$pre
  geom_ribbon(data=newdatYears, aes(ymin = lower, ymax = upper, x = year), fill = "black", alpha=0.1) +
  geom_smooth(data=newdatYears, aes(y = fit, x = year), span = 1, color = "black", size = 1) +
  geom_hline(yintercept=0, linetype="dashed", color = "black") +
    labs(x = "Year of publication",
       y = "Zr") +
  geom_point(aes(x=1990, y= -3), shape = 21, color = "grey65", fill = "grey73", cex = 0.4*15) +
  geom_point(aes(x=1985, y= -3), shape = 21, color = "grey65", fill = "grey73", cex = 0.4*5) +
  annotate("text", x = 1974, y = -3, label = "Precision (1/SE)") +
  annotate("text", x = 1983, y = -3, label = "5") +
  annotate("text", x = 1987, y = -3, label = "15") +
  theme(text =element_text(size = 15))
```

```
plotSmallStudybias / plotYearbias +
  plot_annotation(tag_levels = "A") &
    theme(plot.tag = element_text(face = 'bold'))
```

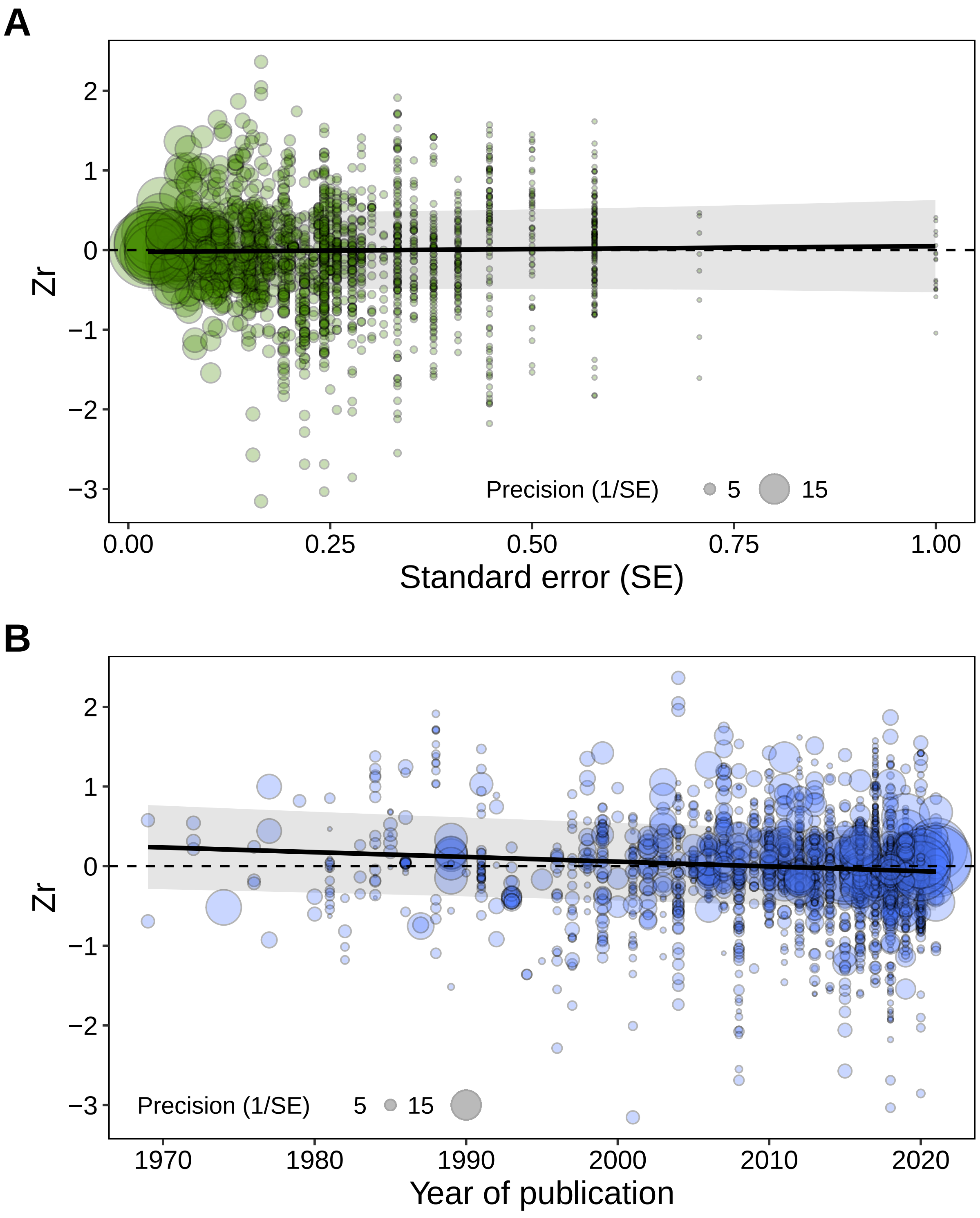


Figure 15: Relationship between A. Standard error and effect size estimates (Zr), and B. Year of publication and effect size estimates (Zr). The points are scaled according to the inverse of their variance, so that larger points are given greater weight in the model and represent more reliable estimates

##### Selection model

```
load(file = "./Models/three.PSM.r0.Rdata")
```

```
plot (three.PSM.r0)
```

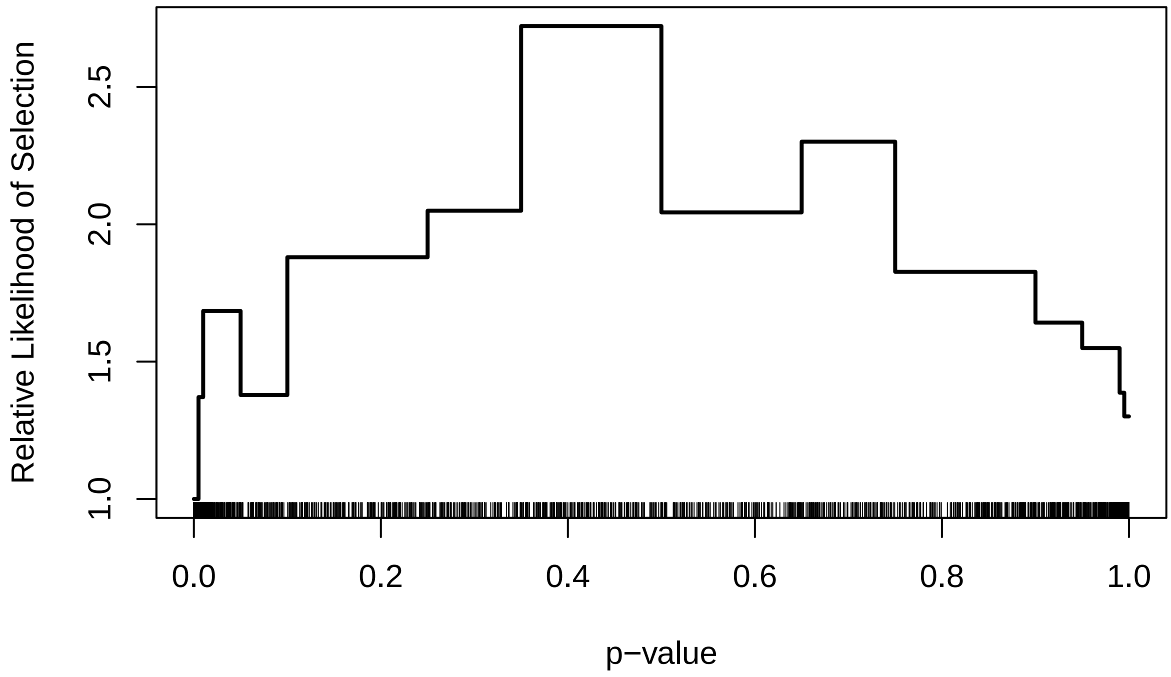


Figure 16: Results of a step function selection model based on several cut-points for a model without any moderators
